# Structural basis for the recognition of K48-linked Ub chain by proteasomal receptor Rpn13

**DOI:** 10.1101/533331

**Authors:** Zhu Liu, Xu Dong, Hua-Wei Yi, Ju Yang, Zhou Gong, Yi Wang, Kan Liu, Wei-Ping Zhang, Chun Tang

## Abstract

The interaction between K48-linked ubiquitin (Ub) chain and Rpn13 is important for proteasomal degradation of ubiquitinated substrate proteins. Only the complex structure between the N-terminal domain of Rpn13 (Rpn13^NTD^) and Ub monomer has been characterized, and it remains unclear how Rpn13 specifically recognizes K48-linked Ub chain. Using single-molecule FRET, here we show that K48-linked diubiquitin (K48-diUb) fluctuates among three distinct conformational states, and a preexisting compact state is selectively enriched by Rpn13^NTD^. The same binding mode is observed for full-length Rpn13 and longer K48-linked Ub chain. Using solution NMR spectroscopy, we have solved the complex structure between Rpn13^NTD^ and K48-diUb. In the structure, Rpn13^NTD^ simultaneously interacts with proximal and distal Ub subunits of K48-diUb that remain associated in the complex, thus corroborating smFRET findings. The proximal Ub interacts with Rpn13^NTD^ similarly as the Ub monomer in the known Rpn13^NTD^:Ub structure, while the distal Ub binds to a largely electrostatic surface of Rpn13^NTD^. Thus, a charge reversal mutation in Rpn13^NTD^ can weaken the interaction between Rpn13 and K48-linked Ub chain, causing accumulation of ubiquitinated proteins. Moreover, blockage of the access of the distal Ub to Rpn13^NTD^ with a proximity attached Ub monomer can also disrupt the interaction between Rpn13 and K48-diUb. Together, the bivalent interaction of K48-linked Ub chain with Rpn13 provides the structural basis for Rpn13 linkage selectivity, which opens a new window for modulating proteasomal function.

## INTRODUCTION

Ubiquitination is a post-translational modification, in which ubiquitin (Ub), a 76-residue protein, is conjugated to a lysine residue of a substrate protein. The modifying Ub is referred to as proximal Ub. Additional Ub units can be further attached to the proximal Ub, in which an isopeptide bond forms between the C-terminal carboxylate group of the incoming Ub (referred to as distal Ub) and an amine group of the proximal Ub. The amine group in the proximal Ub can be one of the seven lysine side chains or N-terminus, for a total of eight homotypic Ub linkages. Ub chains with different linkages encode different cell signals, among which K48-linked Ub chain can target the ubiquitinated substrate protein for proteasomal degradation.

The 26S proteasome is a complex macromolecular machine that is made up of more than 30 types of proteins. Proteasomal degradation can be initiated upon specific interaction between substrate-conjugated K48-linked Ub chain and ubiquitin receptor in the proteasome. Three intrinsic Ub receptors have been identified, including Rpn1^1^, Rpn10/S5a^2^, and Rpn13/Adrm1^3^, while the interactions between ubiquitin-like proteins in shuttle proteins and the proteasome as well as some other mechanisms may also be at play for targeting substrate proteins for degradation^1,4,5^. Structural characterizations of Rpn1, Rpn10, and Rpn13 have shown that, to achieve specific recognition, both proximal Ub and distal Ub in K48-linked diubiquitin (K48-diUb) interact with the two ubiquitin-interacting motifs in Rpn10^2^, or with a toroid of helices in Rpn1^1^. What is puzzling is that, only the complex structure between Ub monomer and Rpn13 N-terminal domain (Rpn13^NTD^) has been determined^6,7^, and only a modest enhancement in binding affinity has been observed for Rpn13 interaction with K48-diUb than with Ub monomer^3^. As a result, it has been previously proposed that only the proximal Ub of K48-diUb interacts with Rpn13, while the distal Ub is free to interact with other proteins^8,9^. Thus, it remains to be established whether and how Rpn13 can interact with two Ub subunits simultaneously in the context of K48-linked Ub chain.

A 26S proteasome comprises a 20S core particle and two 19S regulatory particles stacked on both ends of the 20S particle. Unlike Rpn1 and Rpn10, Rpn13 in the 19S particle has not been visualized from the cryoEM images of human 26S proteasome^10,11^. There are several reasons for the missing electron density of Rpn13. First, biochemical and mass spectrometry analyses showed that only one copy of Rpn13 exists in the 26S proteasome, which means that Rpn13 is at 1:2 molar ratio with other regulatory proteins in the 19S particles^12^. Second, Rpn13 is recruited to the proteasome by interacting with the C-terminal tail of Rpn2^7,13^, which is largely unstructured^11,14^. Therefore, Rpn13 is mobile with respect to the proteasome. Third, Rpn13 itself is dynamic, with a 150-residue long flexible linker connecting Rpn13^NTD^ and the C-terminal domain.

Rpn13 plays redundant roles with Rpn1 and Rpn10 for Ub binding and substrate degradation. Yet evidences have suggested that Rpn13 is uniquely involved in proteasomal degradation of certain proteins. Rpn13 is often over-expressed in malignant diseases^15^. Covalent binding of a small molecule to Rpn13^NTD^, occupying the binding site of Rpn2, can inhibit ER stress-induced protein degradation and induce apoptosis^16,17^. As such, Rpn13 has been identified as an important target for the treatment of multiple myeloma and several other types of cancers. Rpn13 is also responsible for the degradation of IκB for the activation of NF-κB signaling pathway^18^. More interestingly, Rpn13 is subjected to ubiquitination at residue K34^19,20^, and the modification attenuates proteasomal activity by perturbing the interaction between Rpn13 and K48-linked Ub chain^21^. Despite all the biological and pharmaceutical significances of Rpn13, it remains unclear how Rpn13 specifically recognizes K48-linked Ub chain to fulfill its function.

It has been shown that Ub can noncovalently interact with each other, and the noncovalent interactions are modulated by the covalent Ub linkage in di- or poly-Ub chain^22,23^. The ligand free K48-diUb has been crystalized in a fully closed state^24^. However, increasing evidences have indicated that K48-diUb also exists in open conformations in solution^25-27^. Furthermore, it has been shown that ligand-free K48-diUb fluctuates among multiple conformational states, ready to interact with different partner proteins^26,27^. While the open conformational states of K48-diUb are implicated in binding to K48-linkage specific deubiquintase^26^ and to E3-ligase^28^, the fully closed or compact state of K48-diUb is thought to represent a reservoir state^25^. This because in the compact state structure of ligand-free K48-diUb, key hydrophobic residues for interacting with other proteins including L8, I44 and V70, are largely buried at the dimer interface^24,27^, and the two domains may have to open up for binding to specific partner proteins.

To understand how Rpn13 can be specifically recognized by K48-linked Ub chain, we resort to single molecule fluorescence resonance energy transfer (smFRET)^29,30^. The advantage of smFRET is that only two interacting molecules are investigated under near physiological conditions^31^. At the single-molecule level, complications from protein self-association and non-specific interactions, commonly encountered in bulk measurements, can be minimized. Moreover, smFRET measurement allows rapid identification of constituting conformational states of protein structure, which would otherwise be difficult to deconvolute from the ensemble-averaged bulk measurement. In conjunction with smFRET measurements, we have used NMR spectroscopy to determine the solution structure of Rpn13^NTD^:K48-diUb complex. Together, we have shown that K48-diUb alternates among multiple conformational states, and it is the preexisting compact state of K48-linked Ub chain that selectively interacts with Rpn13.

## RESULTS

### K48-linked Ub chain dynamically fluctuates among multiple conformations

We used smFRET to assess the spatial arrangement of the two Ub subunits in ligand-free K48-diUb. We introduced fluorophores, Alexa Fluor 488 and Cy5, at the N-terminus of the distal Ub and the C-terminus of the proximal Ub (Supplementary Fig. S1a). Using expectation maximization algorithm^32^, the smFRET profile of K48-diUb can be best described as three overlapping FRET species (Supplementary Fig. S2). The high-, medium-, and low-FRET species are centered at FRET efficiencies of 0.74, 0.57 and 0.23, with respective populations of ∼48%, ∼39%, and ∼13% (Fig. 1a). The FRET distances between the center of fluorophores are calculated at ∼43 Å, ∼50Å, ∼64 Å, respectively. Thus, the high-, medium-, and low-FRET species can be assigned to open, semi-open, and compact states that are preexisting for K48-diUb.

**Fig. 1.**
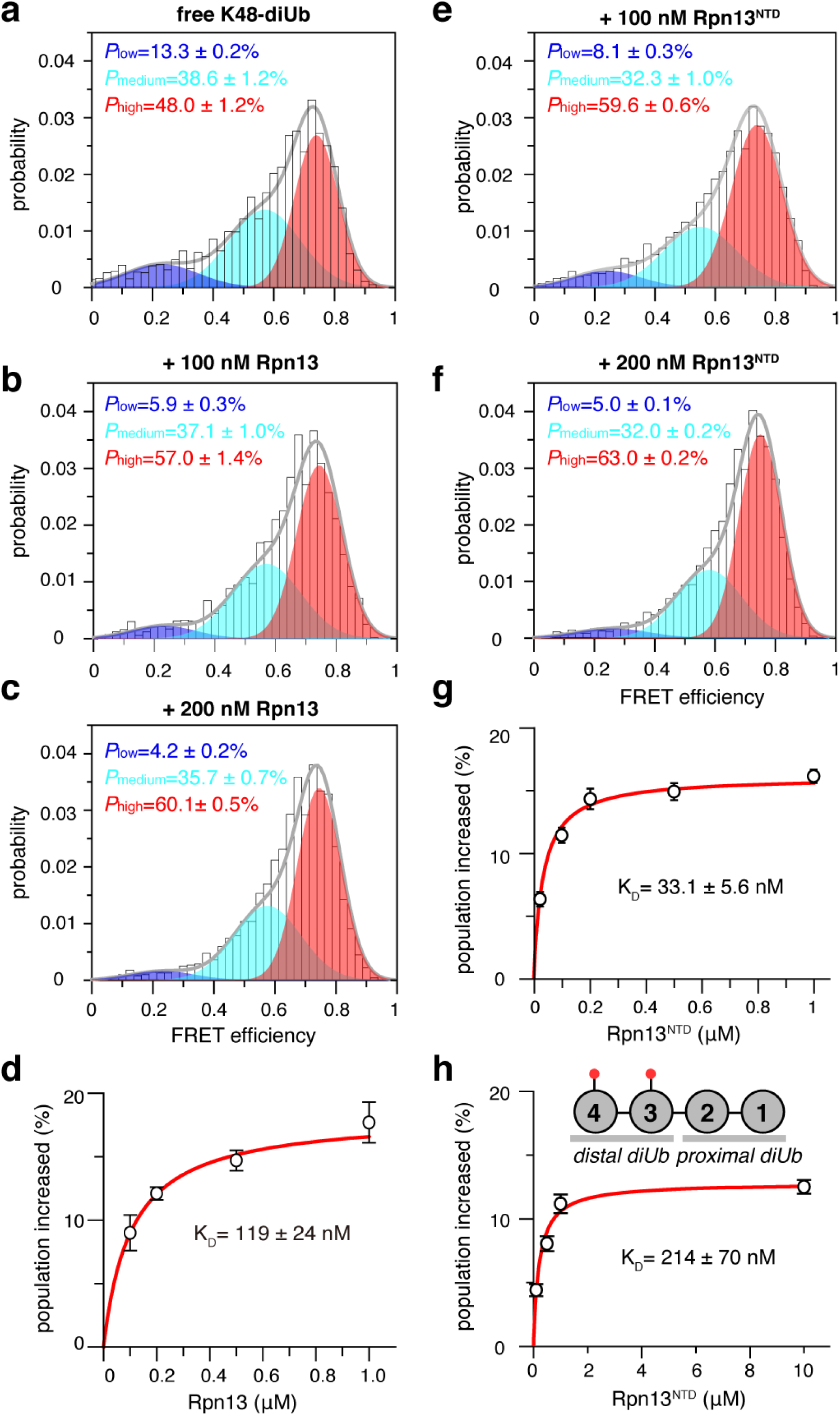
Rpn13 recognizes a preexisting conformational state of K48-diUb. **a** With fluorophores conjugated at 76C site of the proximal Ub and 0C of the distal Ub (cf. Supplementary Fig. S1a), the smFRET profile can be fitted to three overlapping FRET species. Unless otherwise noted, this pair of conjugation sites are used for all smFRET studies of K48-diUb. The high-FRET species (colored red) corresponds to a preexisting compact state. **b-d** The compact state can be selectively enriched by full-length Rpn13, and the population increase can be fitted to a binding isotherm. **e-g** The compact state can be selectively enriched by Rpn13^NTD^, and the population increase can be fitted to be a binding isotherm. **h** With the fluorophores conjugated at N25C/N25C sites, Rpn13^NTD^ can selectively enrich the preexisting compact state of the distal diUb of K48-tetraUb (cf. Supplementary Fig. S5), affording a binding affinity. The populations of the smFRET species are averaged over three independent measurements, with the errors indicating 1 SD; the K_D_ values are reported as best fit ± fitting errors.

The conformational fluctuation of K48-diUb have been previously investigated using smFRET^26^. In that study, the authors resolved two smFRET species for K48-diUb, namely high-FRET and low-FRET species with center FRET efficiencies at 0.69 and 0.41, respectively, in addition to a no-FRET species. The authors showed that the titration of an inactivated OTUB1 (OTUB1i), a deubiquitinase specific for K48 isopeptide linkage^33^, mainly enriches the low-FRET species. Their observation led to the proposal that K48-diUb can be specifically recognized by OTUB1 through a conformational selection mechanism. Here we repeated the smFRET titration, and found that OTUB1i enriches the medium-FRET species (Supplementary Fig. S3a-c). Moreover, the populational increase of the medium-FRET species upon OTUB1i titration can be fitted to a binding isotherm with a K_D_ value of 7.7 ± 0.1 µM (Supplementary Fig. S3d), which is close to the K_D_ value previously reported^26^. Thus, the medium-FRET species in the present study should correspond to the low-FRET species in the previous study, and the discrepancy may arise from different photon counting efficiencies and fitting routines of smFRET time traces.

To further confirm that K48-diUb fluctuates among three preexisting conformational states, we introduced fluorophores at additional pairs of fluorophore conjugation sites (Supplementary Fig. S1b-d). For the alternative sites, although the center efficiencies of the FRET species differ, the smFRET profiles can still be described as three overlapping FRET species with similar populations (Supplementary Fig. S4). For example, for the 25C/25C conjugation sites, the high-, medium-, and low-FRET species are centered at FRET efficiencies of 0.68, 0.54 and 0.21, with respective populations of ∼48%, ∼43%, and ∼9% (Supplementary Fig. S4d). Thus, conjugation of the fluorophores is unlikely to perturb protein structure, and the smFRET measurements have revealed the inherent conformational dynamics of K48-diUb regardless of the conjugation site, that is, the K48-diUb alternates among three distinct states in the absence of a partner protein.

To further assess whether Ub subunits in longer K48-linked Ub chain also fluctuates among multiple conformational states, we analyzed the smFRET profile of K48-linked tetra-ubiquitin (K48-tetraUb). We conjugated the fluorophores at two N25C sites in the distal diUb (with respected to the proximal diUb) of K48-tetraUb. The smFRET profile can also be fit as three overlapping FRET species (Supplementary Fig. S5a). Although the relative populations of the three species are different from that of an isolated K48-diUb with fluorophores conjugated at same sites (Supplementary Fig. S4d), the center FRET efficiencies of the FRET species are almost identical. Thus, the conformational states of the diUb unit is likely preserved in longer K48-linked Ub chain, while the difference in the relative populations can be a result of modulatory effect of the proximal diUb.

### The high-FRET species is selectively enriched by Rpn13

To assess the relationship between the conformational dynamics of K48-linked Ub chain and Rpn13 recognition, we titrated 150 pM fluorophore-labeled K48-diUb with human full-length Rpn13 protein. Interestingly, upon the addition of 100 nM Rpn13, the preexisting high-FRET species of K48-diUb is enriched from ∼48% to ∼57% (Fig. 1a, b), while the population of medium- and low-FRET species decreases. The population of high-FRET species continues to increase with more Rpn13 added (Fig. 1c), and the binding isotherm can be fitted to a K_D_ value of 119 ± 24 nM (Fig. 1d).

It has been previously shown that Rpn13^NTD^ is mainly responsible for Ub binding^6,7^. Thus, we performed smFRET titration for 150 pM fluorophore-labeled K48-diUb using only Rpn13^NTD^ comprising only the first 150 residues. Rpn13^NTD^ also selectively enriches the high-FRET species of K48-diUb (Fig. 1e, f). The population increase of high-FRET species can be fitted to afford a K_D_ value of 33.1 ± 6.9 nM (Fig. 1g), which is about 4-fold increase in affinity as compared to full-length Rpn13. If the fluorophores are attached at alternative conjugation sites (Supplementary Fig. S1b and S4b), titration of Rpn13^NTD^ also causes the enrichment of equivalent FRET species following a similar trend, affording almost identical K_D_ value (Supplementary Fig. S6). Thus, the appearance additional residues may have a small inhibitory effect on the interaction between Rpn13^NTD^ and K48-diUb.

Furthermore, we performed smFRET titration for 150 pM K48-tetraUb with fluorophores conjugated at the distal diUb (Supplementary Fig. S5a). Titration of Rpn13^NTD^ selectively enriches the preexisting high-FRET species of the fluorophore-labeled distal diUb (Supplementary Fig. S5b-d), and the binding isotherm can be fitted to a K_D_ value of 214 ±70 nM (Fig. 1h). The 7-fold reduction in binding affinity compared to Rpn13^NTD^:K48-diUb interaction can be attributed to the self-association between distal diUb and proximal diUb^34^, making the binding surface less available for Rpn13 binding. Importantly, for all the smFRET titrations of K48-diUb or K48-tetraUb, the center efficiency of the high-FRET species changes little in the absence or presence of Rpn13 or Rpn13^NTD^. This means that Rpn13 binds to a preexisting conformation of K48-diUb through a conformational selection mechanism, whether K48-diUb is by itself or part of longer Ub chain. It also means that the high-FRET species, i.e. the preexisting compact state of K48-diUb, is not fully closed, and is ready to interact with other proteins. Importantly, the selective enrichment of the high-FRET species also indicates that Rpn13 should interact with both Ub subunits at the same time.

### Rpn13 preferentially binds to K48-linked diUb

To assess Rpn13 binding specificity for K48 linkage, we assessed the binding affinities between Rpn13 and other types of Ub proteins. With the titration of 200 nM Rpn13^NTD^ into 150 pM fluorophore-labeled K48-diUb, the population of the high-FRET species increases by 15% from ∼48% to ∼63% (Fig. 1f). At this concentration, K48-diUb binding is not yet saturated with Rpn13^NTD^ (Fig. 1g) and therefore, the population of the high-FRET species is sensitive to small change of the available Rpn13^NTD^ concentration. Unlabeled K48-diUb can compete for the binding to Rpn13^NTD^ with fluorophore-labeled K48-diUb. With the addition of 150 pM unlabeled K48-diUb, the population of the high-FRET species decreases by 7.5%, corresponding to a 50% inhibition of Rpn13^NTD^-bound fluorophore-labeled K48-diUb (Fig. 2a). With the addition of 300 pM unlabeled K48-diUb, the population of high-FRET species decreases by 11%, amounting to a total of 73% inhibition (Fig. 2b). As such, both fluorophore-labeled and unlabeled K48-diUb compete for the same binding interface on Rpn13 with similar binding affinity. This also means that conjugation of the fluorophores to K48-diUb causes little perturbation to the interaction between Rpn13^NTD^ and K48-diUb.

**Fig. 2.**
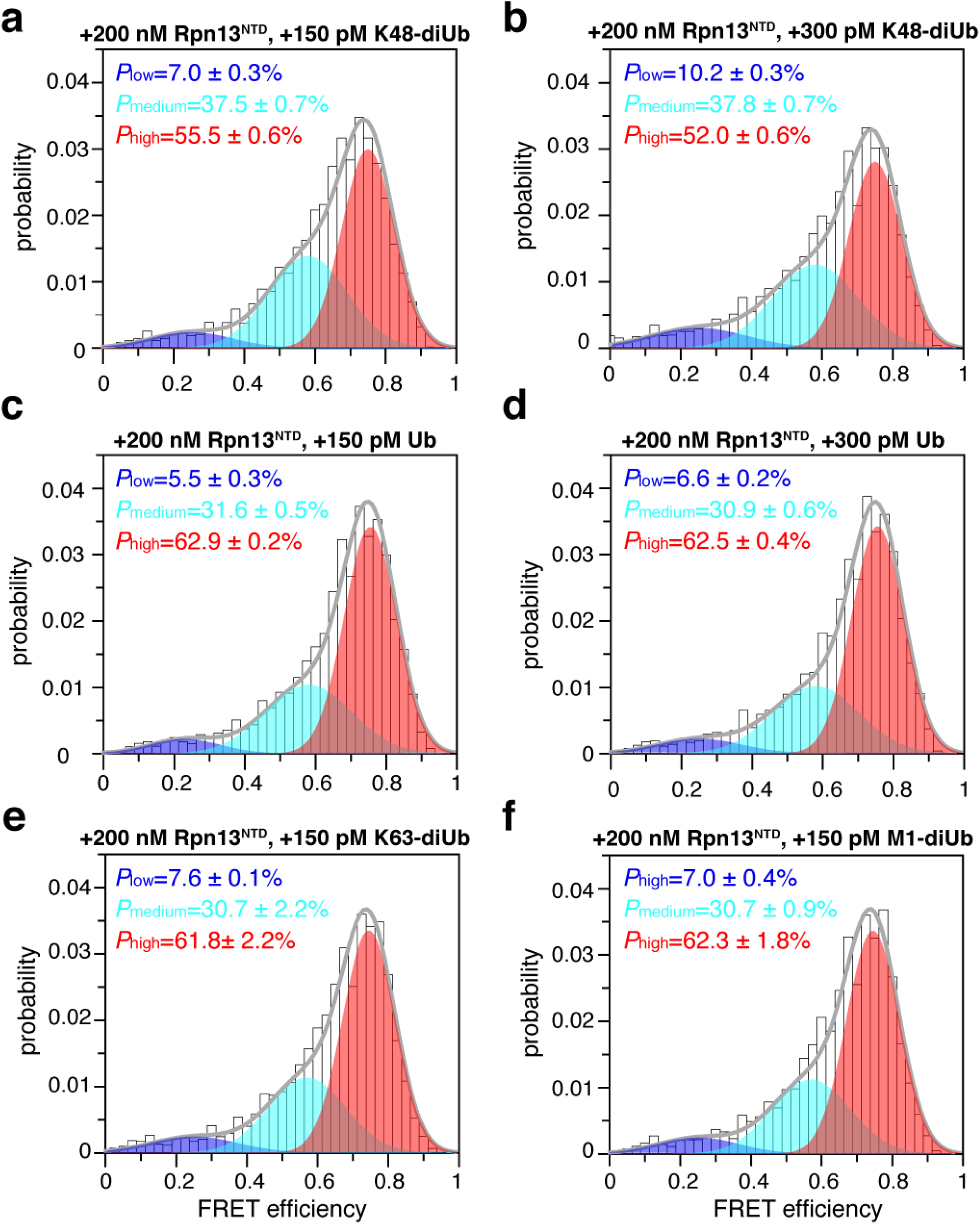
Linkage selectivity of Rpn13. **a,b** Addition of 200 nM Rpn13^NTD^ first enriches the high-FRET species of fluorophore-labeled K48-diUb (cf. Fig. 1f), and further addition of 150 pM and 300 pM unlabeled K48-diUb decrease the population of the high-FRET species. **c,d** Addition of 150 pM and 300 pM unlabeled Ub monomer has negligible effect on the population of the high-FRET species. **e,f** Addition of 150 pM K48-diUb and 150 pM M1-diUb cause very small decrease of the population of the high-FRET species.

Furthermore, to the mixture of 200 nM Rpn13^NTD^ and 150 pM fluorophore-labeled K48-diUb, we added unlabeled Ub monomer, K63-linked diubiquitin or M1-linked diubiquitin, to assess whether other types of Ub can displace K48-diUb. The population of the high-FRET species change little with the addition of 150 pM or 300 pM Ub monomer (Fig. 2c, d). With the addition of 150 pM K63-diUb and 150 pM M1-diUb, the population of the high-FRET species are also unchanged within the error range (Fig. 2e, f). On the other hand, direct titration of 1 µM Rpn13 ^NTD^ into fluorophore conjugated K63-diUb and M1-diUb causes little change to their preexisting smFRET profiles (Supplementary Fig. S7). Together, Rpn13^NTD^ selectively interacts with K48-diUb.

### Solution structure of Rpn13^NTD^:K48-diUb complex

Though we have now shown Rpn13^NTD^ selectively interacts with K48-diUb, only the complex structure between Rpn13^NTD^ and Ub monomer has been determined^6,8^. To understand how the two subunits in K48-diUb can simultaneously interact with Rpn13^NTD^, we set out to determine the solution structure of Rpn13^NTD^:K48-diUb complex using nuclear magnetic resonance (NMR). Upon the formation of protein complex, interfacial residues would experience different local environment and therefore display NMR signals. We found that, titrating unlabeled K48-diUb to ^15^N-labeled Rpn13 causes large chemical shift perturbations (CSPs), which mainly involve residues 73-83 and 93-106 (Fig 3a). These residues form a contiguous surface on Rpn13, covering an area larger than expected from the previously determined complex structure between Rpn13^NTD^ and Ub monomer^6,7^. On the other hand, titrating unlabeled Rpn13^NTD^ to K48-dUb, with either proximal or distal Ub ^15^N-labeled and the other subunit unlabeled, the CSPs are mainly observed for residues in the β-sheet region of both Ub subunits. Some of the interfacial residues also disappeared upon the formation of the complex (Fig 3b, c).

**Fig. 3.**
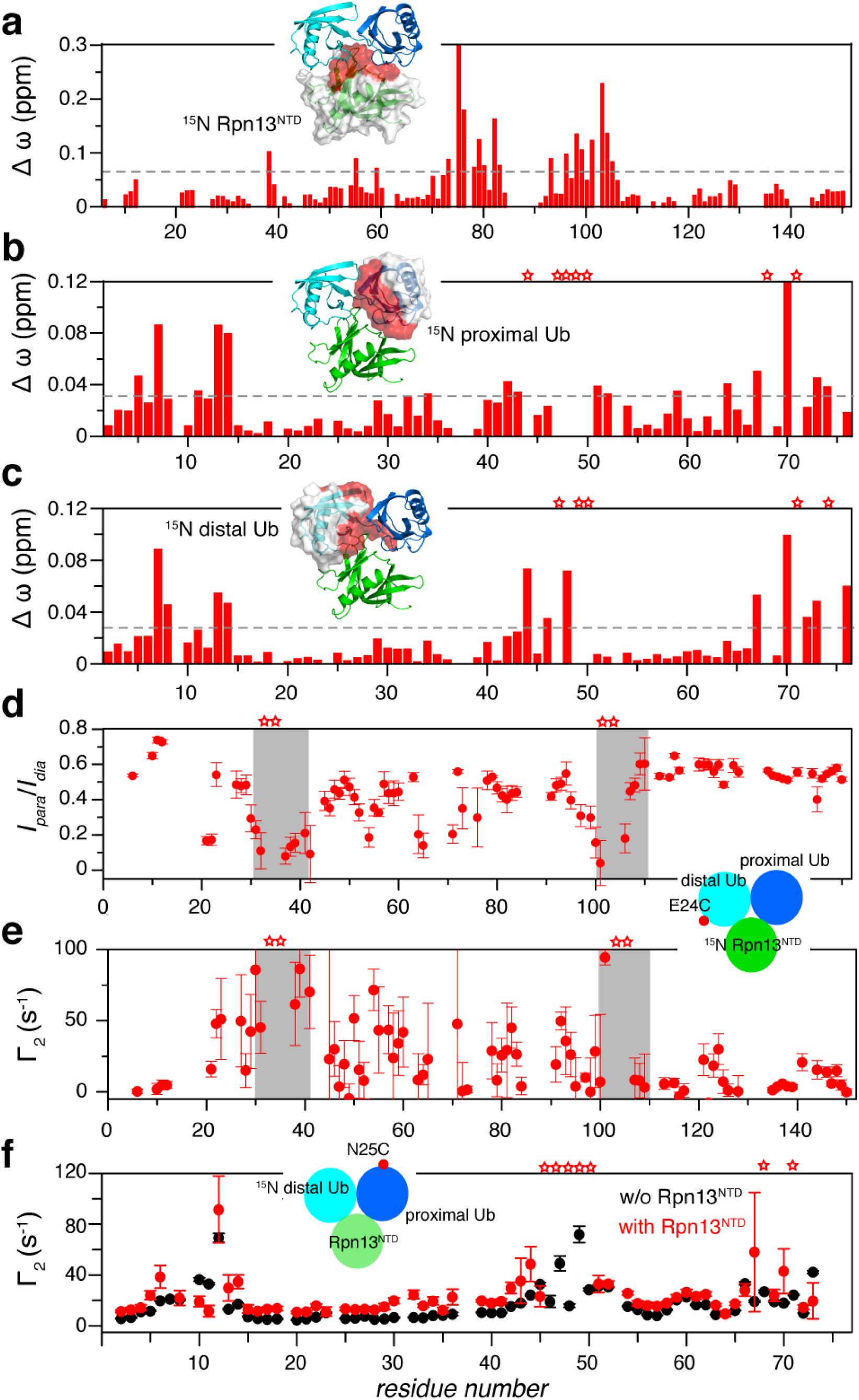
Structural characterization of Rpn13^NTD^:K48-diUb complex using NMR spectroscopy. **a-c** CSP averaged over ^1^H and ^15^N dimensions of backbone amide protons upon the formation of the complex with 0.2 mM isotope-labeled and unlabeled proteins. Insets: the ^15^N-labeled subunit is illustrated with surface representation, and the red colored residues have CSPs above the dotted line. **d,e** With a paramagnetic probe conjugated at E24C site of the distal Ub, the intensity ratios between paramagnetic and diamagnetic spectra, and the PRE Γ_2_ values were assessed for backbone amide protons of ^15^N-labeled Rpn13^NTD^. Gray boxes indicated residues with very large intermolecular PREs. **f** With the paramagnetic probe conjugated at N25C site of the proximal Ub, Γ_2_ values were measured for amide protons of ^15^N-labeled distal Ub in K48-diUb. The error bars indicate 1 S.D., and residues completely broadened out in the complex are denoted with stars.

Nuclear Overhauser effect (NOE) reports distance relationship (< 6 Å) between nuclei. Moreover, ^13^C half-filtered NMR experiment can provide NOE between ^12^C-bonded proton and ^13^C-bonded proton, i.e. intermolecular distance relationship. We obtained intermolecular NOEs between Rpn13^NTD^ and the proximal Ub, between Rpn13^NTD^ and the distal Ub, and between proximal Ub and distal Ub (Supplementary Fig. S8). Furthermore, we conjugated a maleimide-EDTA-Mn^2+^ paramagnetic probe at E24C site of the distal Ub, and collected paramagnetic relaxation enhancement (PRE) for backbone amide protons of Rpn13^NTD^, following the established protocol^22,35^. As assessed either by peak intensity ratio of paramagnetic versus diamagnetic spectra or by transverse relaxation enhancement *Γ*_2_ rate, Rpn13 residues 30-42 and 101-106 experience large PREs with severe line broadening (Fig. 3d, e). We also conjugated the paramagnetic probe at N25C site of the proximal Ub, and evaluated the transverse relaxation enhancement rate Γ_2_ for the distal Ub (Fig. 3f). Large PRE values are observed between the two Ub subunits of the ligand-free K48-diUb, and the addition of Rpn13^NTD^ increases inter-Ub PREs but with a similar PRE profile. The PRE NMR experiments thus confirm that Rpn13^NTD^ enriches the preexisting compact state of K48-diUb.

To refine the Rpn13^NTD^:K48-diUb complex structure against experimental restraints, we performed rigid body docking with torsion angle freedom given to the diUb linker residues and to the side chains of interfacial residues. For the 20 lowest-energy conformers, the root-mean-square (RMS) deviation for backbone heavy atoms of all rigid residues is 0.86 ± 0.54 Å (Supplementary Fig. S9 and Table S1). The two Ub subunits of K48-diUb remain associated in the complex, burying solvent accessible surface area (SASA) of ∼1130 Å^2^. On the other hand, Rpn13^NTD^ wedges in, burying ∼940 Å^2^ of SASA with the proximal Ub and ∼1300 Å^2^ of SASA with the distal Ub (Fig. 4a). The complex structure between Rpn13^NTD^ and proximal Ub of K48-diUb in the present study is similar to the known complex structure between Rpn13^NTD^ and Ub monomer^6,7^, with the RMS difference for backbone heavy atoms 2.17 ± 0.31 Å (Supplementary Fig. S10a). Interestingly, though the hydrophobic residues L8, I44 and V70 in the proximal Ub are involved for interacting with Rpn13, the same three residues in the distal Ub are buried in Ub-Ub interface.

**Fig. 4.**
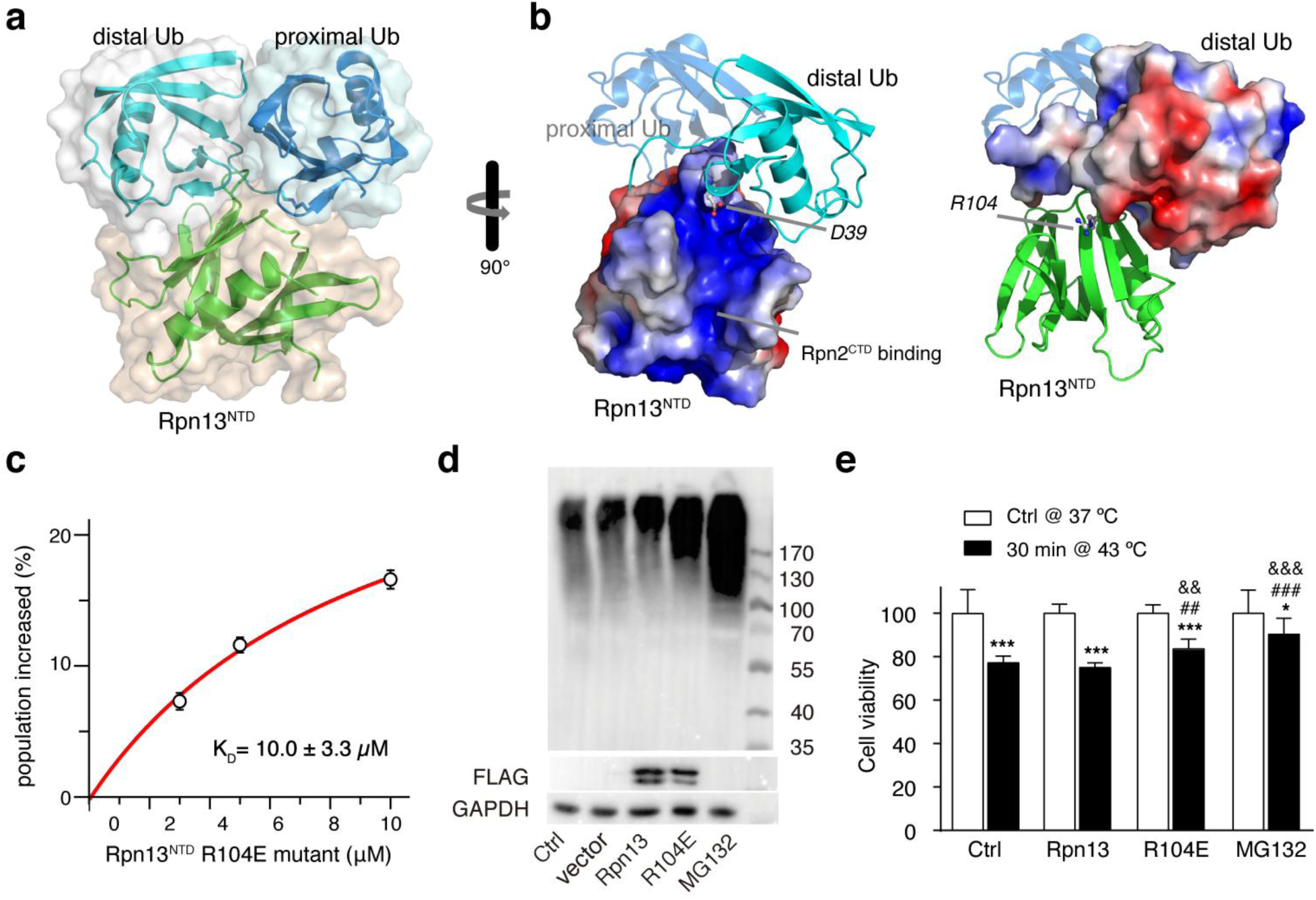
Biological significance for the interaction between Rpn13^NTD^ and the distal Ub of K48-diUb. **a** Structure of Rpn13^NTD^:K48-diUb complex. Formation of the complex buries extensive interfaces between the subunits. **b** Electrostatic potential surfaces of Rpn13^NTD^ and distal Ub of K48-diUb, drawn at ±3 *k*_B_T scale. Residues R104 in Rpn13^NTD^ and D39 in distal Ub are shown as balls-and-sticks. **c** R104E charge-reversal mutation in Rpn13^NTD^ causes a 300-fold decrease in the binding affinity for K48-diUb, as assessed by smFRET (cf. Supplementary Fig. S11). The K_D_ value is reported as best fit ± fitting errors. **d** Western blot analyses show that the transfection of Rpn13 R104E mutant (with an N-terminal flag) increases the overall amount of K48-linked polyUb proteins in cell. **e** Cell viabilities upon heat shock (relative to cells without heat shock) show that transfection of Rpn13 R104E mutant, but not of wildtype Rpn13, confers thermotolerance. \**P*<0.05, \*\*\**P*<0.001, to control cells in the same group, unpaired *t*-test; %%*P*<0.01, ^%%%^*P*<0.001, to the untreated cells with heat shock, one-way ANOVA; ^&&^*P*<0.01, ^&&&^*P*<0.001, to wildtype Rpn13 transfected cells with heat shock, one-way ANOVA.

The Rpn13^NTD^:K48-diUb complex structure can also be corroborated by single-molecule FRET data. Based on the complex structure, we modeled the fluorophores at their conjugation sites in K48-diUb. The average distance is 43.2±5.8 Å between the geometric centers of fluorophore aromatic rings, which corresponds to a theoretical FRET efficiency of 0.73±0.13 (Supplementary Fig. S10b). This value is almost the same as the center efficiency observed for the high-FRET species (Fig. 1a).

### Disruption of Rpn13^NTD^: distal Ub interaction causes accumulation of ubiquitinated proteins in cell

In the complex structure between Rpn13^NTD^ and K48-diUb, the interaction between Rpn13^NTD^ and the proximal Ub is similar to that between Rpn13^NTD^ and Ub monomer, as previously reported (Supplementary Fig. S10a). Thus, we designed experiments to assess functional importance for the interaction between Rpn13^NTD^ and the distal Ub of K48-diUb. Many charged residues are located at the interface between Rpn13^NTD^ and the distal Ub of K48-diUb, and therefore electrostatic force may play an important role for stabilizing the complex (Fig. 4b). Among them, residue D39 in the distal Ub is close to residue R104 in Rpn13. We thus mutated Rpn13 residue R104 to a glutamate, and titrated the mutant Rpn13^NTD^ to fluorophore-labeled K48-diUb. The mutant protein enriches the high-FRET species of K48-diUb (Supplementary Fig. S11). However, the binding affinity becomes much weaker. The binding isotherm can be fitted to a K_D_ value of 10.0 ± 3.3 µM, about 300 times weaker than the wildtype Rpn13^NTD^ (Fig. 4c). As such, the association with the distal Ub is important for the specific recognition between Rpn13 and K48-diUb.

The decreased binding affinity of the R104E mutant allowed us to assess the functional importance of the interaction between Rpn13 and K48-linked Ub chain. Transient transfection of wildtype Rpn13 slightly increases the amount of K48-linked polyUb proteins (Fig. 4d). It is possible that, upon Rpn13 transfection, an excess amount of free Rpn13 compete for binding to the K48-linked polyUb proteins with proteasome-associated Rpn13, thus making the recruitment of ubiquitinated substrate proteins to the proteasome less efficient. On the other hand, the transfection of Rpn13 R104E mutant greatly increases the amount of K48-linked polyUb proteins, as compared to the cells transfected with wildtype Rpn13 (Fig. 4d). As a positive control, we incubated the cells with 1 µM MG132, a potent proteasome inhibitor^36^. Due to the blockage of degradation of labile proteins, addition of MG132 significantly increases the amount of K48-linked polyUb proteins. Taken together, R104E mutation of Rpn13 can lead to the accumulation of ubiquitinated substrate proteins. This can be attributed to weaker interaction between proteasome-associated Rpn13 mutant and K48-diUb and K48-poyUb.

Heat shock can decrease cell viability. We found that 30 min heat shock at 43 °C can decrease the viability of HEK293 cells to 75%. Similar to the previous reports^36,37^, we also found that the treatment of MG132 has a protective effect on cell survival upon heat shock, with cell viability decreased to ∼90% (Fig. 4e). This is because MG132 inhibits proteasomal degradation, making the otherwise short-living heat shock proteins more available (Fig 4d). We also assayed the viability of heat-shocked cells transfected with wildtype Rpn13, and found no significant difference from the control cells without Rpn13 transfection. On the other hand, cell viability of Rpn13 R104E transfected cells decreased to ∼83% upon heat shock, which is significantly higher than that of control cells and cells transfected with wildtype Rpn13 (Fig. 4e). This implies that the mutant Rpn13 has a protective effect on cell survival upon heat shock, similar to the effect of MG132. Taken together, the interaction between Rpn13 and K48-linked Ub chain is essential for Rpn13-mediated recognition of ubiquitinated substrate proteins by the proteasome, while an interfacial point mutation in Rpn13 can cause accumulation of certain substrate proteins such as heat shock proteins, and confer thermotolerance to the transfected cells.

### Rpn13^NTD^:K48-diUb interaction can be targeted to modulate Rpn13 function

Rpn13 is dynamically recruited to the proteasome via the interaction between Rpn13^NTD^ and the C-terminal tail of Rpn2^7,13^. The complex structure here indicates that the binding interface on Rpn13^NTD^ for Rpn2 is close to but does not overlap with the binding interface for the distal Ub of K48-diUb (Fig. 5a). We thus premixed the last 16 residues of Rpn2 (Rpn2^CTD^) with Rpn13^NTD^ or with full-length Rpn13 at 1:1 ratio, and titrated Rpn13^NTD^:Rpn2^CTD^ complex to fluorophore-labeled K48-diUb. The premixing of Rpn2^CTD^ increased the K_D_ value of Rpn13^NTD^:K48-diUb from 33.1 ± 6.9 nM (Fig. 1g) to 66.8 ± 13.9 nM (Supplementary Fig. S12a-c and Fig. 5b), while decreased the K_D_ value of Rpn13:K48-diUb from 119 ± 24 nM (Fig. 1d) to 43.9 ± 12.8 nM (Supplementary Fig. S12d-f and Fig. 5c). Thus, providing the measurement uncertainties, the association of Rpn2^CTD^ cause only small perturbation for the binding affinity between Rpn13 and K48-diUb, which may also have to do with the presence of linker and the C-terminal domain of Rpn13.

**Fig. 5.**
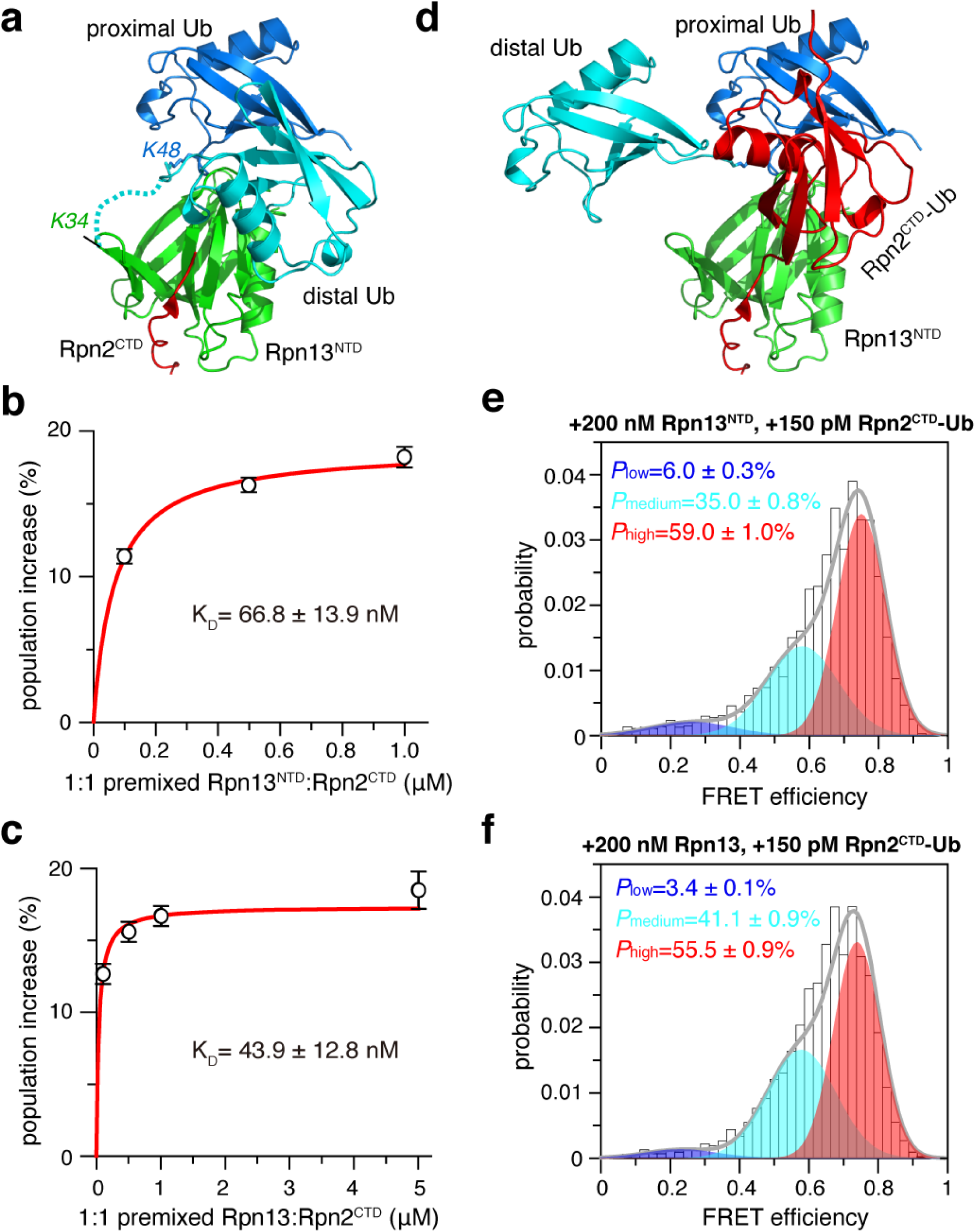
Ub binding surface on Rpn13^NTD^ can be targeted. **a** The binding surface of the distal Ub of K48-diUb on Rpn13^NTD^ is adjacent to the binding surface of Rpn2^CTD^. With Rpn13^NTD^-Rpn2^CTD^ complex structure (PDB code 5V1Y) superimposed by Rpn13^NTD^, Rpn2^CTD^ is shown as red cartoon. Rpn13 residue K34 is close to the C-terminus of distal Ub (shown as a dotted line). **b,c** Binding affinities were obtained from smFRET titrations of equimolar Rpn13^NTD^:Rpn2^CTD^ or Rpn13:Rpn2^CTD^ to fluorophore-labeled K48-diUb. The K_D_ values are reported as best fit ± fitting errors. **d** Binding of Rpn2-anchored Ub monomer likely occupies the same binding surface of the distal Ub, causing the displacement of the latter. **e,f** The smFRET competition experiments, with further addition of 150 pM Rpn2^CTD^-Ub fusion protein, to the mixture of 200 nM Rpn13^NTD^ or full-length Rpn13 and 150 pM fluorophore-labeled K48-diUb (c.f. Fig. 1c and f). The error indicates 1 S.D. from three independent measurements of the populations of the smFRET species.

Rpn2 and distal Ub of K48-diUb occupy nearby surfaces on Rpn13^NTD^. Therefore, a fusion protein with Ub monomer appended at the C-terminus of Rpn2^CTD^ may protrude out and interfere with the interaction between Rpn13^NTD^ with the distal Ub of K48-diUb (Fig. 5d). As we have shown, addition of 200 nM Rpn13^NTD^ increases the population of the high-FRET species of 150 pM fluorophore-labeled K48-diUb to ∼63% (Fig. 1f), while addition of Ub monomer cannot compete for Rpn13NTD binding (Fig. 2c, d). When we added additional 150 pM unlabeled Rpn2^CTD^-Ub fusion protein, the population of high-FRET species decreases by ∼4% to ∼59% (Fig. 5e). On the other hand, we have shown that, addition of 200 nM full-length Rpn13 increases the population of the high-FRET species of 150 pM fluorophore-labeled K48-diUb to ∼60% (Fig. 1c). Further addition of 150 pM unlabeled Rpn2^CTD^-Ub fusion protein decreases the population of high-FRET species by ∼4.5% to ∼55.5% (Fig. 5f). Note that appending a Ub at Rpn2^CTD^ has almost no effect on the interaction between Rpn2^CTD^ and Rpn13^NTD^ (Supplementary Fig. S13). Thus, our data indicate that Rpn2 and K48-diUb binding interfaces on Rpn13^NTD^ are close to each other. More importantly, Rpn2-anchored Ub can physically block the access of the distal diUb to Rpn13, and weaken the interaction between K48-diUb and Rpn13.

## DISCUSSION

In the present study, we have characterized the conformational dynamics of K48-diUb, and elucidated how Rpn13, a proteasomal receptor for ubiquitinated substrate proteins, can specifically interact with K48-linked Ub chain. Our finding thus explains linkage specificity of Rpn13-Ub interaction, and fills a missing piece of the puzzle about proteasomal receptor recognition—specific interactions with Rpn10, Rpn1 and Rpn13 all require the presence of at least two covalently linked Ub subunits. Using the smFRET technique, we show that K48-diUb fluctuates among three conformational states, and the high-FRET species is selectively enriched by Rpn13^NTD^. The selectively also holds for full-length Rpn13 and for K48-trtraUb. The smFRET titrations further indicate that Ub monomer, K63-linked diUb, and M1-linked diUb do not interact with Rpn13. Thus, K48-diUb and K48-polyUb are preferred binding partner of Rpn13.

Using NMR spectroscopy, we have determined the solution structure of the complex between Rpn13^NTD^ and K48-diUb by refining against intermolecular NOE and PRE restraints. The complex structure shows that the interaction between the distal Ub and Rpn13^NTD^ is mainly electrostatic and possibly dynamic, which can explain the sparsity of inter-subunit NOEs between the distal Ub and Rpn13^NTD^. Providing that the distal Ub of K48-diUb transiently opens up, noncovalent interactions can occur between Ub subunits^22^. This would explain a 2:1 stoichiometry between K48-diUb and Rpn13^NTD^, or a 1:1 stoichiometry between Ub monomer and Rnpn13^NTD^ previously observed in bulk titrations^3,7,9^. Indeed, the Rpn13^NTD^-Ub complex has been crystalized as a dimer (Supplementary Fig. S14)^7^. Herein, the structure of Rpn13^NTD^:K48-diUb complex clearly explains how the two Ub subunits of K48-diUb simultaneously interact with Rpn13^NTD^. The NMR structure is also corroborated by the smFRET data, as the calculated and observed FRET efficiencies agree with each other (Supplementary Fig. S9b).

We have also shown that Rpn13^NTD^:K48-diUb interaction can be disrupted by a charge reversal mutation introduced at Rpn13^NTD^ facing the distal Ub of K48-diUb, or by blocking the access of the distal Ub with an Ub monomer fused to Rpn2^CTD^ and anchored at adjacent binding site. Rpn13 itself can be ubiquitinated at residue K34^21^. In Rpn13^NTD^:K48-diUb complex structure, Rpn13 residue K34 is much closer to the C-terminus of the distal Ub of K48-diUb than to the C-terminus of the proximal Ub (Fig. 5a). Owing to high effective concentration, a Ub covalently attached at Rpn13 residue K34 can compete with the distal Ub of K48-diUb for Rpn13 binding, but not with the proximal Ub or Ub monomer as previously proposed^21^. Thus, the present study explains how autoubiquitination can inhibit the function of Rpn13. The binding surface of the distal Ub on Rpn13^NTD^ can also be a novel target of pharmaceutical intervention of proteasomal function.

In summary, we have used smFRET to analyze the number of inter-converting conformational states and their respective populations, and used NMR spectroscopy to determine the structure of the constituting conformational state. Both experiments were conducted in solution, and the FRET data measured for individual molecule and NMR data measured for bulk molecules corroborate each other. As such, the conjoint use of bulk and single-molecule measurements gives us better insight of protein dynamics and the mechanism of protein function. We envision such integrative use of the two solution-based techniques can be generally used for characterizing other multi-subunit and multi-domain proteins.

## MATERIALS AND METHODS

### Protein sample preparation

Human ubiquitin was cloned to pET-11a vector (Novagen), and point mutations were introduced using the QuikChange strategy using Kod plus enzyme (Toyobo). To ensure a single final product of diubiquitin, K48R (or K63R) mutation was introduced to the distal Ub and 77D was introduced to the proximal Ub. For site-specific conjugation of fluorescent probes (or paramagnetic probe), cysteine mutations were included to ubiquitin, one at a time, including 76C (mutated from Gly76), 25C (mutated from Asn25), 24C (mutated from Glu24), and 0C (MSAC sequence appended at the N-terminus). Human Rpn13^NTD^ (residues 1-150) was cloned into a pET-11a vector. With a C91A mutation introduced to human OTUB1 (residues 40-271) and with a hexa-histidine tag appended at protein N-terminus, the resulting protein, OTUB1i, has no deubiquitinase activity and was also cloned to pET-11a vector. Rpn2^CTD^ (GPQEPEPPEPFEYIDD in sequence at the very C-terminus of Rpn2) was purchased from GL Biochem (Shanghai). Rpn2^CTD^ was also appended at the N-terminus of ubiquitin. The fusion protein was purified in the same way as Ub monomer, using Sepharose Q, Source-Q and Sephacryl S100 columns (GE Healthcare) sequentially.

Ubiquitin monomer, M1-diUb, or Rpn2^CTD^-Ub was expressed in BL21 star cells; the expression was induced with the addition of 1 mM IPTG at OD_600_ value of ∼0.8, and the culture was continued to grow at 37 °C for ∼4 hours. Expression of Rpn13^NTD^ or full-length Rpn13 was induced with the addition of 0.2 mM IPTG at OD_600_ of 0.8, and the culture was gown overnight at 20 °C. Expression of OTUB1i was induced with the addition of 50 µM IPTG at OD_600_ of 0.8, and the culture was gown overnight at 16 °C. For expressing isotopically labeled protein, U-^15^N-labeled NH_4_Cl and U-^13^C-labeled glucose (Sigma-Aldrich) were used as the sole nitrogen and carbon sources, and the cells were grown in M9-minimum medium prepared either H_2_O or D_2_O (Sigma-Aldrich). Rpn13^NTD^, ubiquitin (or ubiquitin mutants), or M1-diUb was purified on Sepharose SP, Sephacryl S100 and Source-S columns. OTUB1 was purified from His-affinity and Superdex-75 columns. The full length Rpn13 was purified with sequential use His-Trap, Source-Q and Sephacryl S100 columns (GE Healthcare). Rpn2^CTD^-Ub was purified using Sepharose Q, Source-Q and Sephacryl S100 sequentially. The identities of the proteins were confirmed using electrospray ionization mass spectrometry (Bruker Daltonics), and the protein concentrations were measured based on UV absorptions at 280 nm.

The ligation between two ubiquitin proteins to afford a diubiquitin was performed following the established protocol^38^. To ensure a unique ligation product, 0.5 mM proximal Ub appended with residue D77 at the C-terminus was mixed equimolar with the distal Ub carrying a K48R mutation in pH 8.0 buffer with 10 mM Tris•HCl and 1mM DTT. A mixture of 2.5 μM E1, 20 μM E2-25K, 2 mM ATP, 5 mM MgCl_2_, 10 mM creatine phosphate (Sigma-Aldrich, catalogue number 27920), 2 U/ml inorganic pyrophosphatase (Sigma-Aldrich, catalogue number I1643), and 1U/ml creatine phosphokinase (Sigma-Aldrich, catalogue number C3755) were added for the reaction. The ligation took place at 30 °C for 6 hours and was quenched with the addition of 5 mM DTT and 5 mM EDTA. The K48-diUb protein was purified from the reaction mixture using Superdex 75 and Source-S columns (GE Healthcare) in tandem. To prepare K48-diUb with site-specific cysteine mutations, mutant proteins were used as the starting material for enzyme-catalyzed ligation reaction. Two copy of human ubiquitin genes were fused together to generate M1-linked diubiquitin (M1-diUb). M1-diUb protein was prepared in the same way as Ub monomer. K63-linked diubiquitin (K63-diUb) was prepared as described previously^23^.

We used Alexa Fluor 488 C_5_ maleimide (Thermo-Fisher, catalogue number A10254) as the fluorescence donor and Cy5 maleimide (GE Healthcare, catalogue number PA15131) as the fluorescence acceptor. The Förster distance R_0_ between the dyes is 52 Å^39^. The dyes were first dissolved in DMSO at 1 mM concentration, and were freshly prepared just before conjugation. The DTT in the buffer for cysteine mutant of K48-diUb was removed by desalting (HiPrep 26/10 Desalting column, GE Healthcare), and the protein was reacted with the pre-mixed excess Alexa488 and Cy5 dyes. The conjugation reaction was performed in dark at room temperature for 4 hours, and the product was purified from the reaction mixture using Source-Q column (GE Healthcare). The doubly labeled K48-diUb had absorptions at 280 nm, 493 nm and 650 nm all three wavelengths, and was further confirmed using mass spectrometry (Bruker Daltonics). Similarly, mixed labeling of fluorophores were introduced at 76C site of the proximal Ub and 25C site of the distal Ub in K63-diUb and M1-diUb.

To prepare K48-linked tetraUb, residue D77 of the proximal Ub in K48-diUb was deblocked using yeast ubiquitin C-terminal hydrolase-1 (YUH1), which was then ligated to another Ub77D protein, thus to generate tri-ubiquitin (triUb). Subsequently, residue D77 in the most proximal Ub of triUb was deblocked and was ligated to a fourth Ub77D protein to generate tetraUb. To prepare K48-tetraUb with cysteine mutations at specific sites, the mutant proteins were used as the starting material for enzyme-catalyzed ligation reactions. Fluorophore conjugation was performed as described above for K48-diUb.

### Single-molecule FRET data collection and analysis

A Nikon A1 confocal microscope was used for single molecule imaging, and a pulsed interleaved excitation (PIE) scheme^40^ was employed, by using two SPCM-AQRH detectors (Excelitas) to record the fluorescence time traces at the emission wavelengths of donor and acceptor. Two picosecond pulsed diode laser heads (LDH-P-C-485B and LDH-P-C-640B, PicoQuant) were driven by a PDL 828 Sepia II driver (PicoQuant) at 32 MHz repetition rate and were used for the interleaved excitations of Alexa488 and Cy5. Each laser was coupled to the A1 microscope through a single-mode fiber and was reflected by a dichroic mirror through a water immersion objective (WI 60×, NA 1.20, Nikon).

The protein sample was loaded to a hybridization chamber (Thermo Fisher, catalogue number S24732). The fluorescence outputs were recorded with TimeHarp 260 PCI board (PicoQuant) built into a PC workstation, and the data were stored in the time-tagged time-resolved module (TTTR, PicoQuant). The photon counts for donor and acceptor emissions were analyzed by bursts^41,42^ using the parameters previously described^43^.

The smFRET measurements were performed at 25 °C in 20 mM pH 7.4 HEPES buffer containing 150 mM NaCl, 0.01% (v/v) Tween 20 (Thermo Fisher), 1 mM ascorbic acid and 1 mM methylviologen (Sigma-Aldrich). The concentration of the doubly labeled K48-diUb was ∼150 pM. All other unlabeled protein samples were all prepared as stock solutions in 20 mM pH 7.4 HEPES buffer containing 150 mM NaCl and 4 mM DTT and were mixed with K48-diUb prior to each experiment to the final desired concentration. The time traces were binned with 1-ms bins and the threshold for photon count traces was typically 3-5 counts/bin per burst depending on the background dark counts. With high concentrations of ubiquitin receptors added, the threshold for photon count could reach 10 counts/bin, owing to higher background fluorescence.

To determine the number of constituting conformational states for K48-diUb from the smFRET data, we used expectation maximization (EM) algorithm^32^ for fitting multi-Gaussian mixture model using an in-house program. First, the program makes the hypothesis about the number of different components that make up the observed smFRET data, and then it calculates the likelihood for the corresponding parameters, and finally the program adjusts the parameters to maximize the likelihood. For evaluating the number of Gaussian components, we introduced Bayesian information criterion (BIC) and Akaike information criterion (AIC)^44,45^, in addition to the correlation coefficient *R*^2^. The smFRET measurement and fitting were repeated three times, and the averaged FRET efficiencies, populations and associated standard deviations were reported. To estimate the K_D_ value, the populational increase of a particular smFRET species was fitted against the concentration of unlabeled protein added in Origin 8.0, using the following equation:

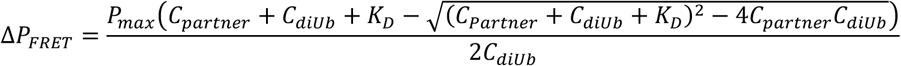

Here Δ*P*_FRET_ is the change of the population of a particular FRET species, *P*_max_ is the maximum difference for the FRET species with or without the partner protein, *C*_partner_ is the concentration of the partner protein added, *C*_diUb_ is the concentration of the fluorophore-conjugated K48-diUb.

### NMR data acquisition and analysis

For the assignment of NMR resonances of Rpn13^NTD^, U-[^2^H, ^13^C, ^15^N]- and U-[^13^C, ^15^N]-labeled proteins were prepared in 20 mM pH 6.5 MES buffer containing 150 mM NaCl, 4 mM DTT, and 10% D_2_O. The NMR data were collected for a ∼0.6 mM sample at 25 °C on a Bruker Avance III 850 MHz spectrometer equipped with a TXI cryogenic probe. The ^1^H-^15^N HSQC, HNCA, HNCB and HCCH-TOCSY experiments were used for backbone and side-chain resonance assignment; the final assignment was compared against the known assignment^13^.

To characterize the interactions between Rpn13^NTD^ and K48-diUb, K48-diUb with the proximal Ub 15N-labeled was titrated with unlabeled Rpn13^NTD^, K48-diUb with the distal Ub ^15^N-labeled was titrated with unlabeled Rpn13^NTD^, or ^15^N-labeled Rpn13^NTD^ was titrated with unlabeled K48-diUb. Average CSP for each residue was calculated using the equation Δδ=[0.5(Δδ_H_^2^+0.2Δδ_N_^2^)]^1/2^, in which Δδ_H_ and Δδ_N_ are the CSP values in ^1^H and ^15^N dimensions. Both K48-diUb and Rpn13^NTD^ are 0.2 mM in concentration, and the 1:1 complex contains ^15^N-labeled and unlabeled proteins.

To obtain intermolecular NOEs, we prepared two NMR samples. In one sample, the distal Ub is Ub U-[^13^C,^15^N]-labeled, and the proximal Ub and Rpn13^NTD^ are unlabeled. In the other sample, Rpn13^NTD^ is U-[^13^C,^15^N]-labeled, and K48-diUb is unlabeled. The concentrations were 0.5 mM for the isotopically labeled protein, and 0.6 mM for the unlabeled protein. Half-filtered ^13^C NOESY experiments^46^ were performed with a 120 ms NOE mixing time. Intermolecular NOE cross-peaks were assigned manually and iteratively based on the initial structures for the ternary complex.

Cysteine point mutation was introduced either at the proximal Ub (N25C site) or distal Ub (E24C site), and the mutant protein with natural isotope abundance was conjugated to maleimide-EDTA-Mn^2+^ paramagnetic probe as previously described^22,23^. The conjugated product was further purified with Source-Q column, and was confirmed by mass spectrometry. The paramagnetically tagged Ub is enzymatically ligated to another Ub with either ^15^N enrichment (N25C site) or with natural isotope abundance (E24C site). Transverse relaxation rates Γ_2_ were measured for the backbone amide protons of 0.3 mM K48-diUb using the established protocol^47^ in the absence or presence of equimolar Rpn13^NTD^.

### Structure calculation of the Rpn13^NTD^:K48-diUb complex

Structural calculation was performed using Xplor-NIH^48^ with conjoined torsion angle/rigid body simulated annealing refinement^49^. The starting structures of ubiquitin and Rpn13^NTD^ were taken from PDB structures 1UBQ^50^ and 2R2Y^6^, respectively. The initial structure of Rpn13^NTD^ was further refined using intramolecular NOE and residual dipolar coupling restraints collected in pf1 and PEG-hexanol media. The coordinates for ubiquitin were duplicated, and an isopeptide bond was patched between the proximal Ub and distal Ub. The refinement was performed with a target function comprising the intermolecular NOE restraints (between proximal Ub and distal Ub, between proximal Ub and Rpn13^NTD^, and between distal Ub and Rpn13^NTD^), a van der Waals repulsive term, and a radius-of-gyration restraint^51^ for the interfacial residues mapped by the NMR chemical shift perturbation, and the PRE-restraints with the paramagnetic probe patched at N25C site of proximal Ub and at E24C of distal Ub (applied as squared-well potential with ± 10 s^-1^ upper/lower bounds).

To obtain an initial pose for the ternary complex, for residues 1-71 of the proximal Ub, residues 1-71 of the distal Ub and Rpn13^NTD^, each was grouped as a rigid body, while complete torsion angle freedom was given to the isopeptide linker including the side chain of K48 in the proximal Ub and residues 72-76 in the distal Ub. Besides experimental restraints, the diubiquitin linker residues were subjected to torsion angle database restraint^52^ for better packing at the interfaces. The solvent accessible surface area (SASA) was calculated with Xplor-NIH with a probe radius of 1.4 Å, and was subtracted from the SASA calculated from the sub of surface area of the free proteins to obtain the buried SASA. Based on the complex structure, fluorescent probes (maleimide AlexaFluor-488 and Cy5) were patched onto the labeling sites, and the linker between protein backbone and rigid portion of the fluorophore (also including residues 72-76 when labeled at the Ub C-terminus) are given torsion angle freedom and are allowed reorient. The distances between the fluorophores are calculated with <*r*^-6^> averaging to afford expected FRET efficiencies. Structures were selected for their overall energy, and figures were illustrated using PyMol version 2.0 (Schrödinger LLC).

### Cell-based experiments

Human Rpn13 cDNA was purchased (Sino Biologicals, Beijing, China), and was sub-cloned to a pcDNA3.1 vector. A flag tag was appended at the N-terminus of Rpn13, and R104E mutation was site-specifically introduced. The plasmids (pcDNA3.1, pcDNA3.1/flag-hRpn13, and pcDNA3.1/flag-hRpn13/R104E) were transfected to HEK293 cells (Cell Biology of the Chinese of Academy of Sciences, Shanghai, China) using PEI reagent (polyethyleneimine, m.w. 25,000, Alfa Aesar Chemical, Shanghai, China). As a positive control, 6 hours after transfection, MG132 (at 1 μM final concentration, Sigma-Aldrich, St. Louis MO, USA) was added to one group of un-transfected cells. 48 hours after transfection, the cells were washed twice with ice-cold phosphate-buffered saline (PBS) and then lysed at 4°C in RIPA lysis buffer (Beyotime Biotechnology, Shanghai, China). Protein concentrations were determined by using BCA assay kit (Beyotime Biotechnology).

For Western blotting, 50 μg cell lysate was separated in 7.5% SDS page gel, and transferred to 0.45 μm PVDF membrane (Millipore Corporation, Billerica, USA). The membrane was blocked by 5% fat-free milk, and was incubated with rabbit anti-K48 linkage specific polyubiquitin antibody (Cell Signaling Technology, Boston, USA), rabbit anti-flag tag antibody (Proteintech, Wuhan, China) and mouse anti GAPDH antibody (Proteintech) at 4°C overnight. After being washed three time by using TBST (Tris buffered saline with Tween 20, 50 mM Tri-HCl, 0.15 M sodium chloride, 0.05% Tween 20, pH 7.6), the membrane was incubated in horse HRP-conjugated anti-mouse IgG antibody (Cell Signaling Technology) or goat HRP-conjugated anti-rabbit IgG antibody (Cell Signaling Technology) at room temperature for 2 hours. After being washed three time with TBST, the membrane was incubated with ECL (Enhanced Chemiluminescence, Bio-RAD, Hercules, USA), and the immunoblots were analyzed by ChemiDocTM MP V3 (Bio-RAD).

As a parallel experiment, 6 hours after transfection, the three groups of transfected cells and two groups of un-transfected cells were cultured in two 96-well plates (with 1 µM MG132 added to one group of un-transfected cells). 24 hours later, the cells in one 96-well plate was switched to 43 °C culture temperature using pre-heated culture medium, and were cultured at 43 °C for 30 min. The heat-shocked cells were reversed to 37 °C, and were cultured for another 24 hours. The cells in the other 96-well plate was cultured under normal conditions at 37 °C. MTT assay was used to detect the cell viability; cells were incubated with 0.5 mg/ml 3-(4,5-cimethylthiazol-2-yl)-2,5-diphenyl tetrazolium bromide (MTT, Sigma-Aldrich) for 2 hours at 37 °C. After the removal of the MTT solution, 100 μl DMSO were added. Following another 10-minute incubation, the absorbance was read at 570 nm in a plate reader (Elx800, Bi-TEK Instrument USA). Cell viabilities were normalized to those of 37 °C control cells.

### Accession codes

Atomic coordinates for complex structure between K48-diUb and Rpn13 N-terminal domain have been deposited with the PDB with accession code of 5YMY and the associated NMR data have been deposited at BMRB with the accession code of 36127.

## ACKNOWLEDGMENTS

The work has been supported by the National Key R&D Program of China (2018YFA0507700 to C.T., W.P.Z., Z.L. and Z.G, 2016YFA0501200 to C.T., Z.G. and X.D., and 2017YFA0505400 to X.D.) and by the National Natural Science Foundation of China (91753132 and 31770799 to C.T., 81573400 to W.P.Z., 31500595 to Z.L., 31400735 to Z.G., and 31400644 to X.D.).

## AUTHOR CONTRIBUTIONS

Z.L., W.-P.Z. and C.T. designed the experiments, Z.L., H.-W.Y. and J.Y. prepared the samples, Z.L., H.-W.Y. and K.L. performed smFRET experiments and analyses, J.Y. and Y.W. performed cell-based experiments, X.D. and Z.G. determined the complex structure, W.-P.Z. and C.T. wrote the manuscript with support from all authors.

## REFERENCES

1. Shi, Y. et al. Rpn1 provides adjacent receptor sites for substrate binding and deubiquitination by the proteasome. Science 351(2016).

2. Deveraux, Q., Ustrell, V., Pickart, C. & Rechsteiner, M. A 26-S Protease Subunit That Binds Ubiquitin Conjugates. J. Biol. Chem. 269, 7059–7061 (1994).

3. Husnjak, K. et al. Proteasome subunit Rpn13 is a novel ubiquitin receptor. Nature 453, 481–488 (2008).

4. Hjerpe, R. et al. UBQLN2 Mediates Autophagy-Independent Protein Aggregate Clearance by the Proteasome. Cell 166, 935–949 (2016).

5. Samant, R.S., Livingston, C.M., Sontag, E.M. & Frydman, J. Distinct proteostasis circuits cooperate in nuclear and cytoplasmic protein quality control. Nature 563, 407–411 (2018).

6. Schreiner, P. et al. Ubiquitin docking at the proteasome through a novel pleckstrin-homology domain interaction. Nature 453, 548–552 (2008).

7. VanderLinden, R.T., Hemmis, C.W., Yao, T., Robinson, H. & Hill, C.P. Structure and energetics of pairwise interactions between proteasome subunits RPN2, RPN13, and ubiquitin clarify a substrate recruitment mechanism. J. Biol. Chem. 292, 9493–9504 (2017).

8. Zhang, N. et al. Structure of the s5a:k48-linked diubiquitin complex and its interactions with rpn13. Mol. Cell 35, 280–290 (2009).

9. Chen, X. et al. Structures of Rpn1 T1:Rad23 and hRpn13:hPLIC2 Reveal Distinct Binding Mechanisms between Substrate Receptors and Shuttle Factors of the Proteasome. Structure 24, 1257–1270 (2016).

10. Huang, X., Luan, B., Wu, J. & Shi, Y. An atomic structure of the human 26S proteasome. Nat. Struct. Mol. Biol. 23, 778–785 (2016).

11. Chen, S. et al. Structural basis for dynamic regulation of the human 26S proteasome. Proc. Natl. Acad. Sci. U. S. A. 113, 12991–12996 (2016).

12. Berko, D. et al. Inherent asymmetry in the 26S proteasome is defined by the ubiquitin receptor RPN13. J. Biol. Chem. 289, 5609–5618 (2014).

13. Lu, X. et al. Structure of the Rpn13-Rpn2 complex provides insights for Rpn13 and Uch37 as anticancer targets. Nat. Commun. 8, 15540 (2017).

14. Wang, X. et al. Molecular Details Underlying Dynamic Structures and Regulation of the Human 26S Proteasome. Mol. Cell. Proteomics 16, 840–854 (2017).

15. Pilarsky, C., Wenzig, M., Specht, T., Saeger, H.D. & Grutzmann, R. Identification and validation of commonly overexpressed genes in solid tumors by comparison of microarray data. Neoplasia 6, 744–750 (2004).

16. Anchoori, R.K. et al. A bis-benzylidine piperidone targeting proteasome ubiquitin receptor RPN13/ADRM1 as a therapy for cancer. Cancer Cell 24, 791–805 (2013).

17. Trader, D.J., Simanski, S. & Kodadek, T. A reversible and highly selective inhibitor of the proteasomal ubiquitin receptor rpn13 is toxic to multiple myeloma cells. J. Am. Chem. Soc. 137, 6312–6319 (2015).

18. Mazumdar, T. et al. Regulation of NF-kappa B activity and inducible nitric oxide synthase by regulatory particle non-ATPase subunit 13 (Rpn13). Proc. Natl. Acad. Sci. U. S. A. 107, 13854–13859 2010).

19. Wagner, S.A. et al. A proteome-wide, quantitative survey of in vivo ubiquitylation sites reveals widespread regulatory roles. Mol. Cell. Proteomics 10, M111 013284 (2011).

20. Kim, W. et al. Systematic and quantitative assessment of the ubiquitin-modified proteome. Mol. Cell 44, 325–340 (2011).

21. Besche, H.C. et al. Autoubiquitination of the 26S proteasome on Rpn13 regulates breakdown of ubiquitin conjugates. EMBO J. 33, 1159–1176 (2014).

22. Liu, Z. et al. Noncovalent dimerization of ubiquitin. Angew. Chem. Int. Ed. 51, 469–472 (2012).

23. Liu, Z. et al. Lys63-linked ubiquitin chain adopts multiple conformational states for specific target recognition. eLife 4, e05767 (2015).

24. Cook, W.J., Jeffrey, L.C., Carson, M., Chen, Z. & Pickart, C.M. Structure of a diubiquitin conjugate and a model for interaction with ubiquitin conjugating enzyme (E2). J. Biol. Chem. 267, 16467–16471 (1992).

25. Hirano, T. et al. Conformational dynamics of wild-type Lys-48-linked diubiquitin in solution. J. Biol. Chem. 286, 37496–37502 (2011).

26. Ye, Y. et al. Ubiquitin chain conformation regulates recognition and activity of interacting proteins. Nature 492, 266–270 (2012).

27. Berlin, K. et al. Recovering a Representative Conformational Ensemble from Underdetermined Macromolecular Structural Data. J. Am. Chem. Soc. 135, 16595–16609 (2013).

28. Liu, S. et al. Promiscuous interactions of gp78 E3 ligase CUE domain with polyubiquitin chains. Structure 20, 2138–2150 (2012).

29. Schuler, B. Single-molecule FRET of protein structure and dynamics - a primer. J. Nanobiotechnology 11 Suppl 1, S2 (2013).

30. Dimura, M. et al. Quantitative FRET studies and integrative modeling unravel the structure and dynamics of biomolecular systems. Curr. Opin. Struct. Biol. 40, 163–185 (2016).

31. Lerner, E. et al. Toward dynamic structural biology: Two decades of single-molecule Forster resonance energy transfer. Science 359(2018).

32. Burnham, K.P. Multimodel Inference: Understanding AIC and BIC in Model Selection. Soci. Meth. Res. 33, 261–304 (2004).

33. Wiener, R., Zhang, X., Wang, T. & Wolberger, C. The mechanism of OTUB1-mediated inhibition of ubiquitination. Nature 483, 618–622 (2012).

34. Satoh, T. et al. Crystal structure of cyclic Lys48-linked tetraubiquitin. Biochem. Biophys. Res. Commun. 400, 329–333 (2010).

35. Liu, Z., Gong, Z., Dong, X. & Tang, C. chim. Biophys. Acta, Proteins Proteomics 1864, 115–122 (2016).

36. Lee, D.H. & Goldberg, A.L. Proteasome inhibitors cause induction of heat shock proteins and trehalose, which together confer thermotolerance in Saccharomyces cerevisiae. Mol. Cell. Biol. 18, 30–38 (1998).

37. Kim, H.J. et al. Systemic analysis of heat shock response induced by heat shock and a proteasome inhibitor MG132. PloS one 6, e20252 (2011).

38. Pickart, C.M. & Raasi, S. Controlled synthesis of polyubiquitin chains. Methods Enzymol. 399, 21–36 (2005).

39. Kalinin, S. et al. A toolkit and benchmark study for FRET-restrained high-precision structural modeling. Nat. Methods 9, 1218–1225 (2012).

40. Muller, B.K., Zaychikov, E., Brauchle, C. & Lamb, D.C. Pulsed interleaved excitation. Biophys. J. 89, 3508–3522 (2005).

41. Lee, N.K. et al. Accurate FRET measurements within single diffusing biomolecules using alternating-laser excitation. Biophys. J. 88, 2939–2953 (2005).

42. Gopich, I.V. Accuracy of maximum likelihood estimates of a two-state model in single-molecule FRET. J. Chem. Phys. 142(2015).

43. Dong, X. et al. Ubiquitin S65 phosphorylation engenders a pH-sensitive conformational switch. Proc. Natl. Acad. Sci. U. S. A. 114, 6770–6775 (2017).

44. Akaike, H. A new look at the statistical model identification. IEEE Trans. Autom. Control 19, 716–723 (1974).

45. Schwarz, G. Estimating the dimension of a model. Ann. Stat. 6, 461–464 (1978).

46. Zwahlen, C. et al. Methods for measurement of intermolecular NOEs by multinuclear NMR spectroscopy: Application to a bacteriophage lambda N-peptide/boxB RNA complex. J. Am. Chem. Soc. 119, 6711–6721 (1997).

47. Iwahara, J., Tang, C. & Clore, G.M. Practical aspects of 1H transverse paramagnetic relaxation enhancement measurements on macromolecules. J. Magn. Reson. 184, 185–195 (2007).

48. Schwieters, C.D., Kuszewski, J.J. & Clore, G.M. Using Xplor-NIH for NMR molecular structure determination. Prog. Nucl. Magn. Reson. Spectrosc. 48, 47–62 (2006).

49. Clore, G.M. Accurate and rapid docking of protein-protein complexes on the basis of intermolecular nuclear overhauser enhancement data and dipolar couplings by rigid body minimization. Proc. Natl. Acad. Sci. U. S. A. 97, 9021–9025 (2000).

50. Vijay-Kumar, S., Bugg, C.E. & Cook, W.J. Structure of ubiquitin refined at 1.8 A resolution. J. Mol. Biol. 194, 531–544 (1987).

51. Tang, C. & Clore, G.M. A simple and reliable approach to docking protein-protein complexes from very sparse NOE-derived intermolecular distance restraints. J. Biomol. NMR 36, 37–44 (2006).

52. Bermejo, G.A., Clore, G.M. & Schwieters, C.D. Smooth statistical torsion angle potential derived from a large conformational database via adaptive kernel density estimation improves the quality of NMR protein structures. Protein Sci. 21, 1824–1836 (2012).

