## Supplemental figure S1- S14 and supplemental table S1 for "Structural basis for the recognition of K48-linked Ub chain by proteasomal receptor Rpn13"

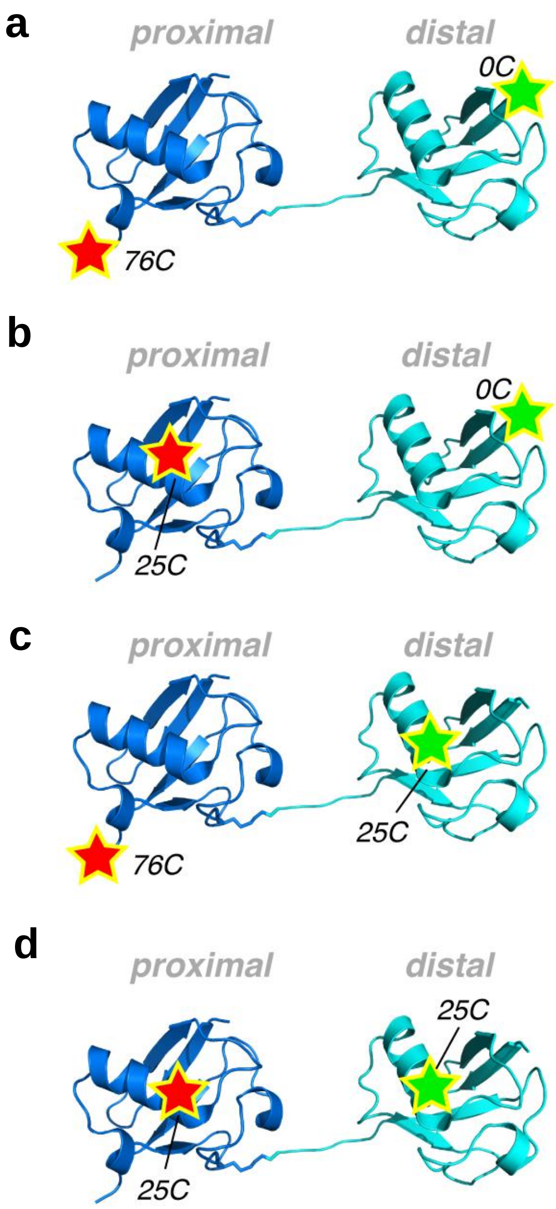

**Supplementary Fig. S1** Pairs of fluorophore conjugation sites used in the present study. Cysteine point mutations were introduced to the C-terminus (76C site, mutated from Gly76), the N-terminus (0C site with MSAC appended), or 25C site (mutated from Asn25). With Alexa Fluor 488 and Cy5 dyes conjugated, four different K48-diUb samples were obtained for smFRET measurement. The proximal Ub here is colored blue, and the distal Ub is colored cyan.

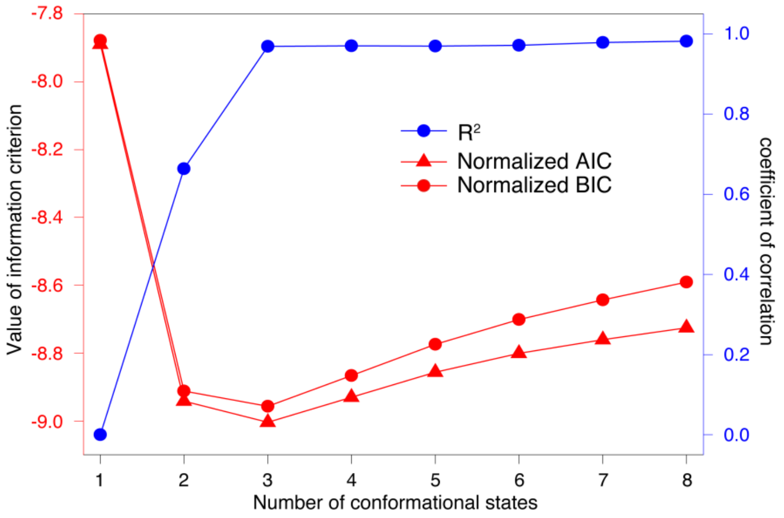

**Supplementary Fig. S2** The smFRET data of K48-diUb can be best fitted as three overlapping FRET species. We used AIC and BIC criteria to identify the optimal number of conformational states. Shown here is the analysis for the smFRET data obtained with the fluorophores attached 76C site of the proximal Ub and 0C of the distal Ub of K48-diUb (c.f. Supplementary Fig. S1a). In addition to AIC and BIC criteria, the overall correlation coefficient $R^2$ was assessed, which is defined as 1-RSS/TSS (RSS, sum of square of residuals; TSS, total sum of squares). Note that $R^2$ value increases to 0.962 with a three-state model, while the normalized AIC and BIC values also achieve minima.

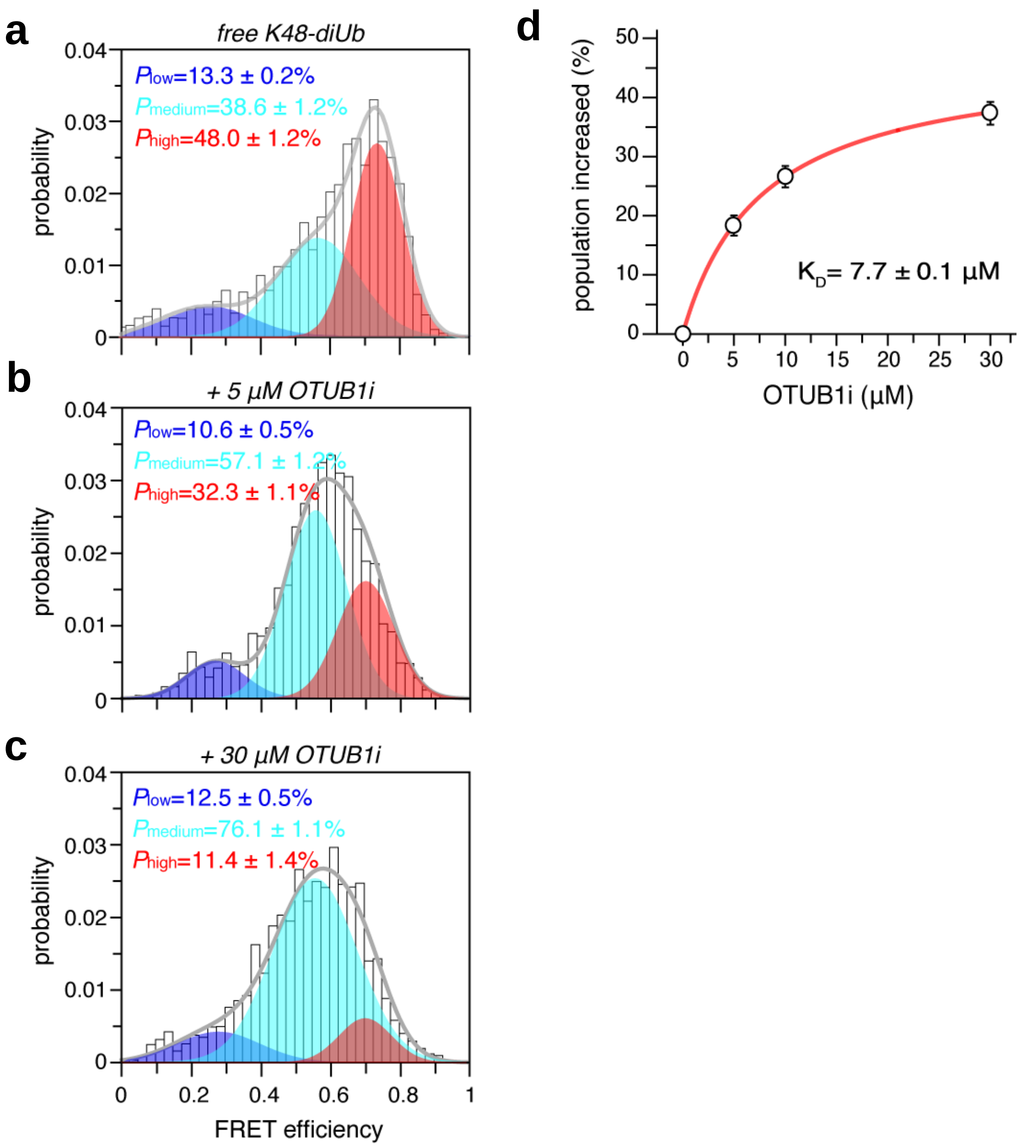

**Supplementary Fig. S3** Titration of OTUB1i increases the population of the medium FRET species. (**a**-**c**) OTUB1 is a deubiquitinase specific for K48-linkage, while OTUB1i carries an active site mutation, C91A. Fluorophores (Alexa Fluor 488 and Cy5) were introduced at 76C site of the proximal Ub and 0C of the distal Ub (Supplementary Fig. S1a). Titration of OTUB1i specifically enriches the medium FRET species, while leaving the centers of the three FRET species unperturbed. (**d**) The population increase of the medium FRET species is plotted vs. the concentration of OTUB1i. The curve can be fitted to a binding isotherm with a $K_D$ value of ~7.7 µM. Averaged populations and standard deviations were obtained from three independent measurements of smFRET time traces.

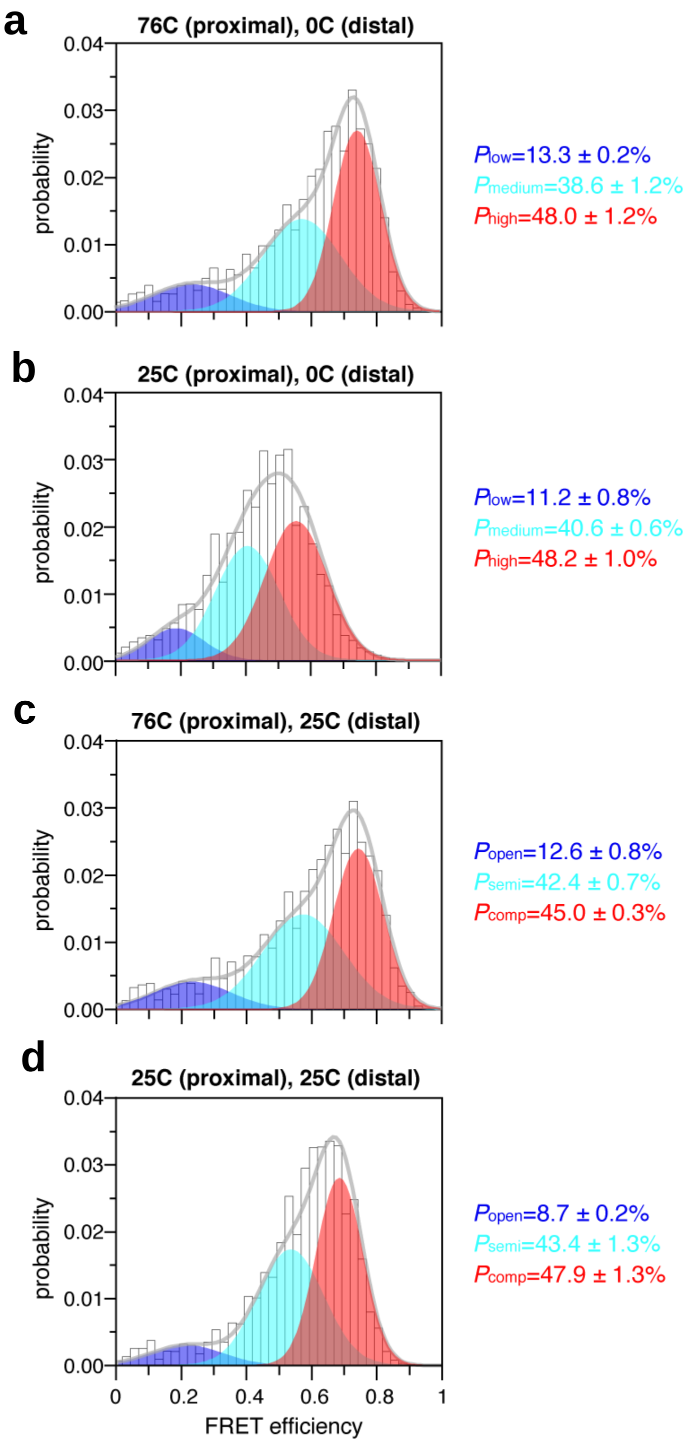

**Supplementary Fig. S4** Fluorophore labeling does not perturb the intrinsic dynamics of K48-diUb. Regardless of the conjugation sites (cf. Supplementary Fig. S1), the smFRET data can be described as the sum of three overlapping FRET species. Though the population of each FRET species is about the same, that the center of the species is different due to the different inter-dye distances. Averaged populations and standard deviations were obtained from three independent measurements of smFRET time traces.

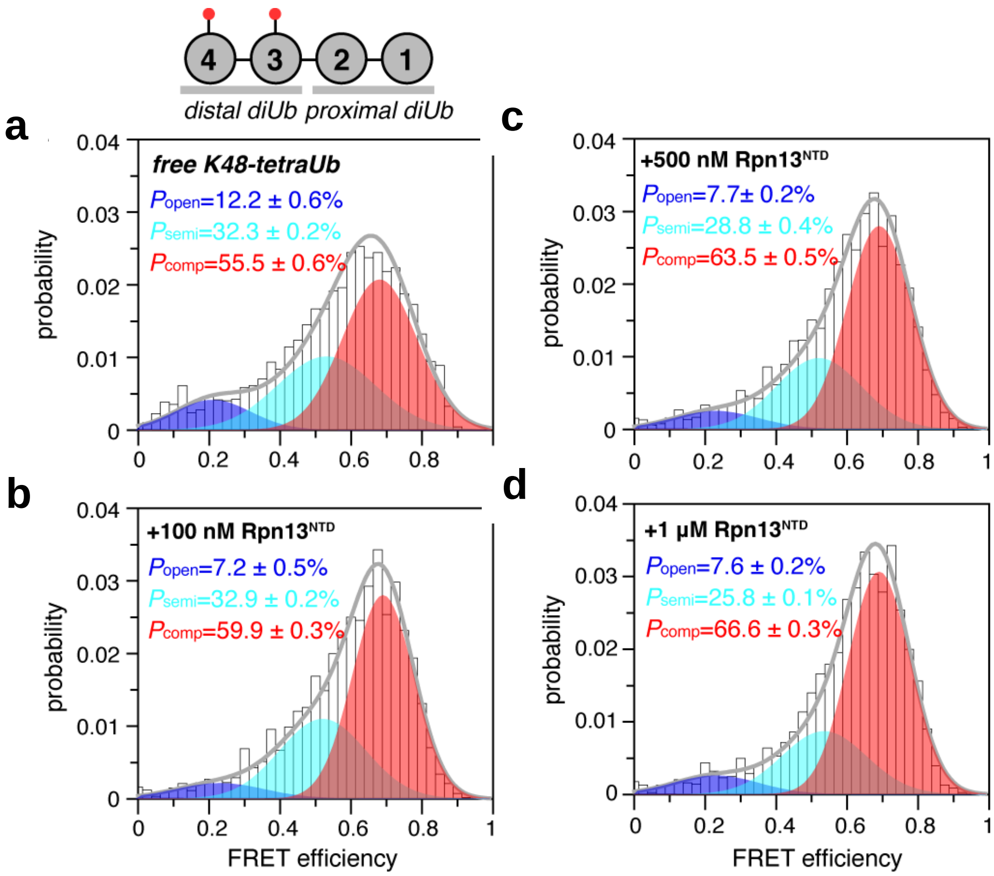

**Supplementary Fig. S5** The smFRET titration of Rpn13$^{NTD}$ into K48-tetraUb. Fluorophores are conjugated at N25C/N25C sites of the distal diUb in K48-tetraUb (cf. Supplementary Fig. S1d). The conformational space of the distal diUb appears similar to an isolated K48-diUb, alternating among thee conformational states. Increasing concentrations of Rpn13$^{NTD}$ selectively enriches the high-FRET species that corresponds to a preexisting compact conformational state. Averaged populations and standard deviations were obtained from three independent measurements of smFRET time traces.

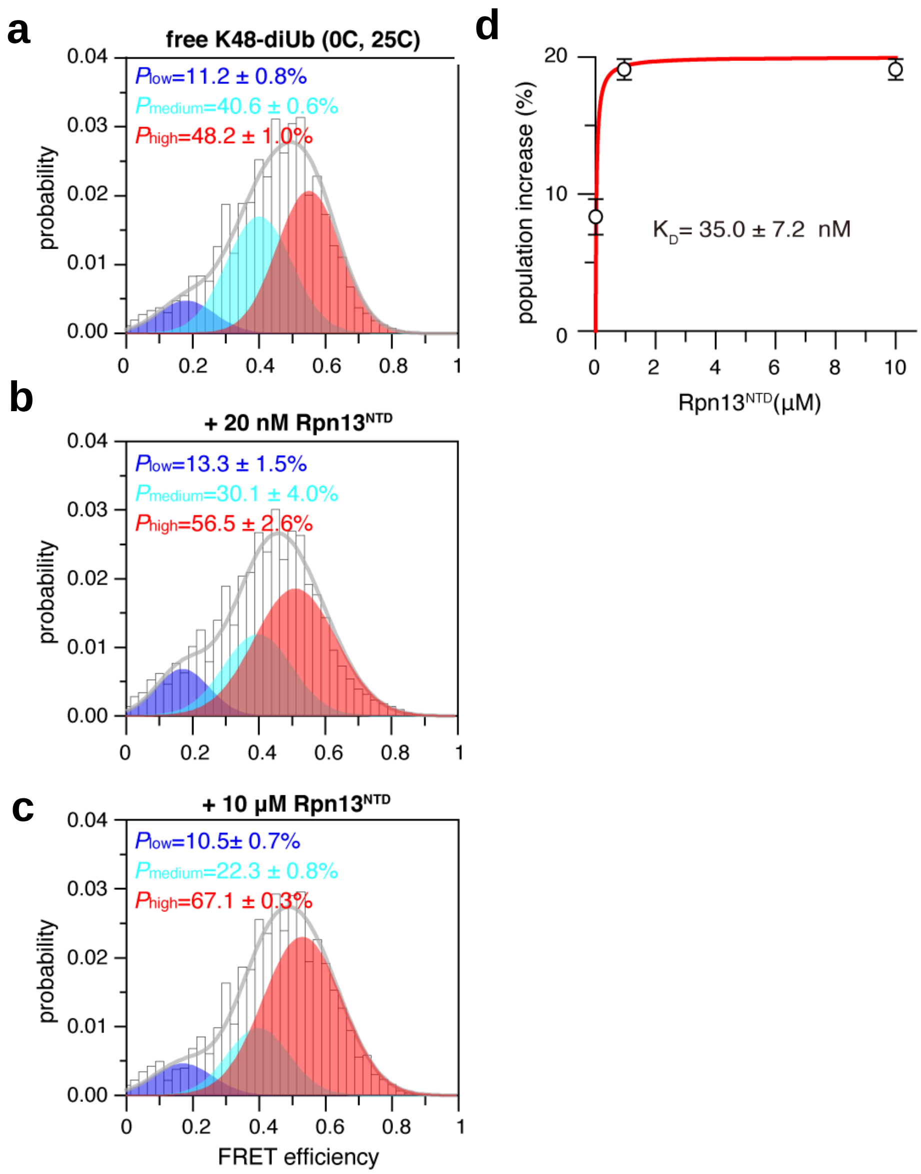

**Supplementary Fig. S6** The smFRET titration of K48-diUb with Rpn13$^{NTD}$, with the fluorophores labeled at alternative pair of residues. Rpn13$^{NTD}$ selectively enriches the high FRET species, with fluorophores (Alexa Fluor 488 and Cy5) introduced at 25C site of the proximal Ub and 0C of the distal Ub (cf. Supplementary Fig. S1b and Fig. S4b). At 20 nM and 10 μM, the high-FRET species is enriched by ~8% and ~19%, respectively. The population increase can be fitted to a binding isotherm. Averaged populations and standard deviations were obtained from three independent measurements of smFRET time traces; $K_D$ value is reported as best fit ± fitting error.

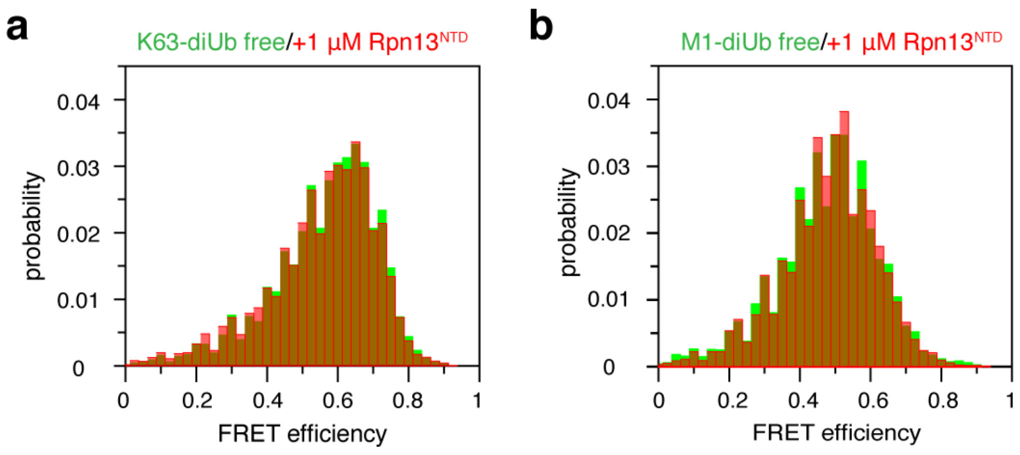

**Supplementary Fig. S7** The smFRET profiles for (**a**) K63-diUb and (**b**) M1-diUb. Alexa488 and Cy5 are conjugated at 76C site of the proximal Ub and 25C site of the distal Ub. The smFRET profiles of free diUb proteins are shown as transparent green; with the addition of 1 μM $Rpn13^{NTD}$, the smFRET profiles are shown as transparent red.

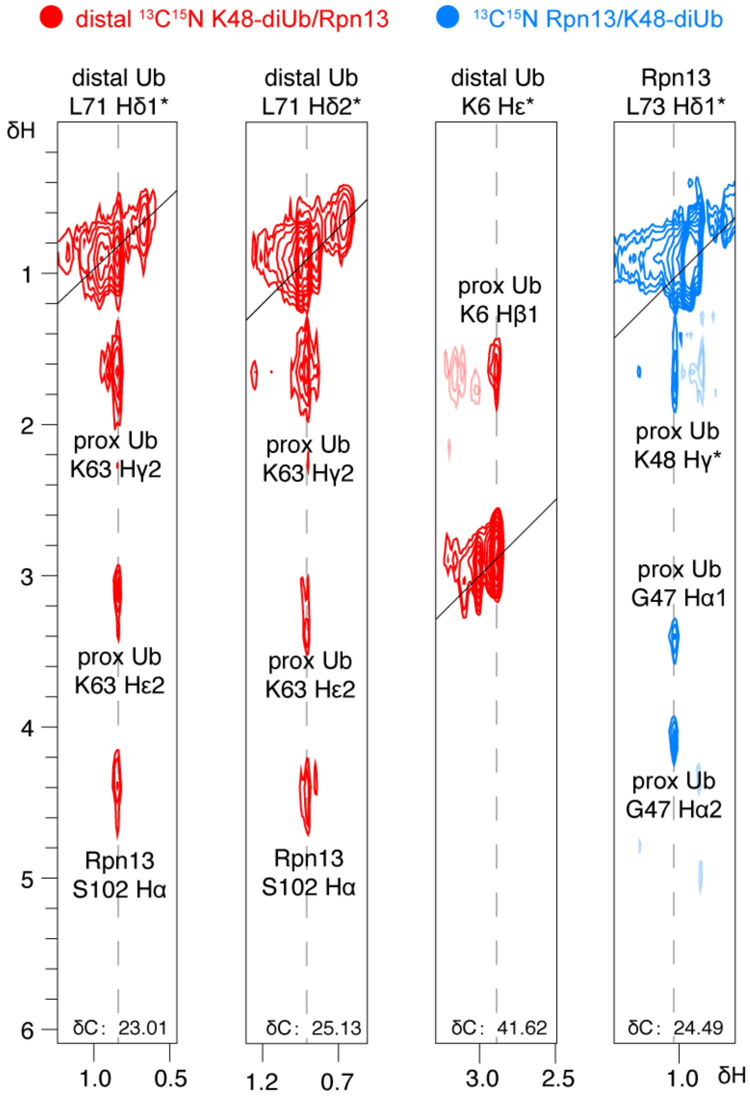

**Supplementary Fig. S8** Representative strips showing intermolecular NOEs between Rpn13$^{NTD}$ and K48-diUb. The sample was prepared with distal Ub of K48-diUb U- [$^{13}$C,$^{15}$N]-labeled and Rpn13$^{NTD}$ unlabeled, or with Rpn13$^{NTD}$ U- [$^{13}$C,$^{15}$N]-labeled and K48-diUb unlabeled. For the half-filtered 3D-NOESY experiment, the residues and atoms to which the NOE cross-peaks are correlated are labeled. The chemical shift in ppm for the corresponding $^{13}$C planes are labeled at the bottom of each strip.

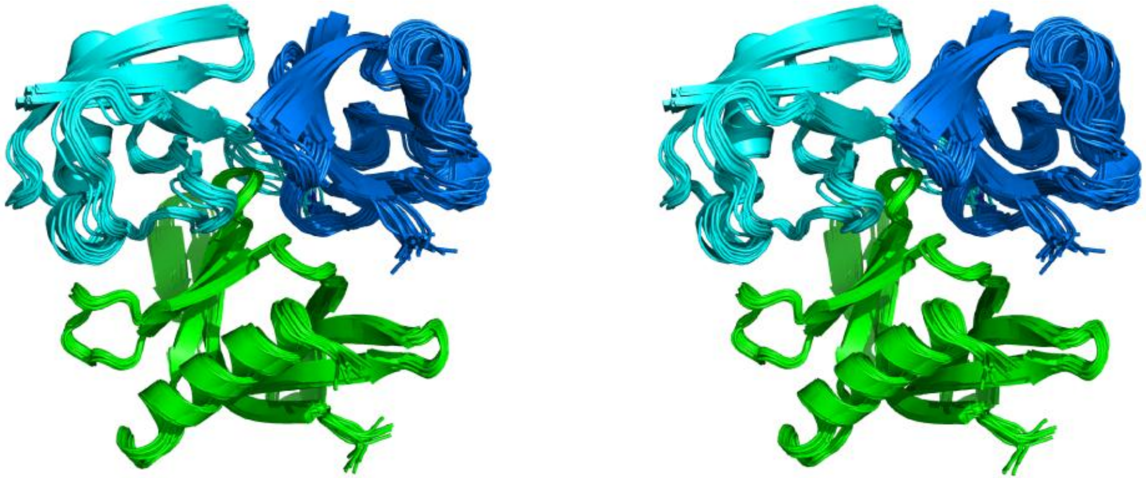

**Supplementary Fig. S9** Solution structure of Rpn13$^{NTD}$:K48-diUb complex. Shown in stereoscopic view, a total of 20 conformers were superimposed. Rpn13$^{NTD}$, proximal Ub and distal Ub are colored green, blue and cyan, respectively. The conformers were selected from 128 calculated for their lowest overall energy and root-mean-square (RMS) deviations from the mean. The overall RMS deviation for the backbone heavy atoms of these is 0.86 ±0.54 Å.

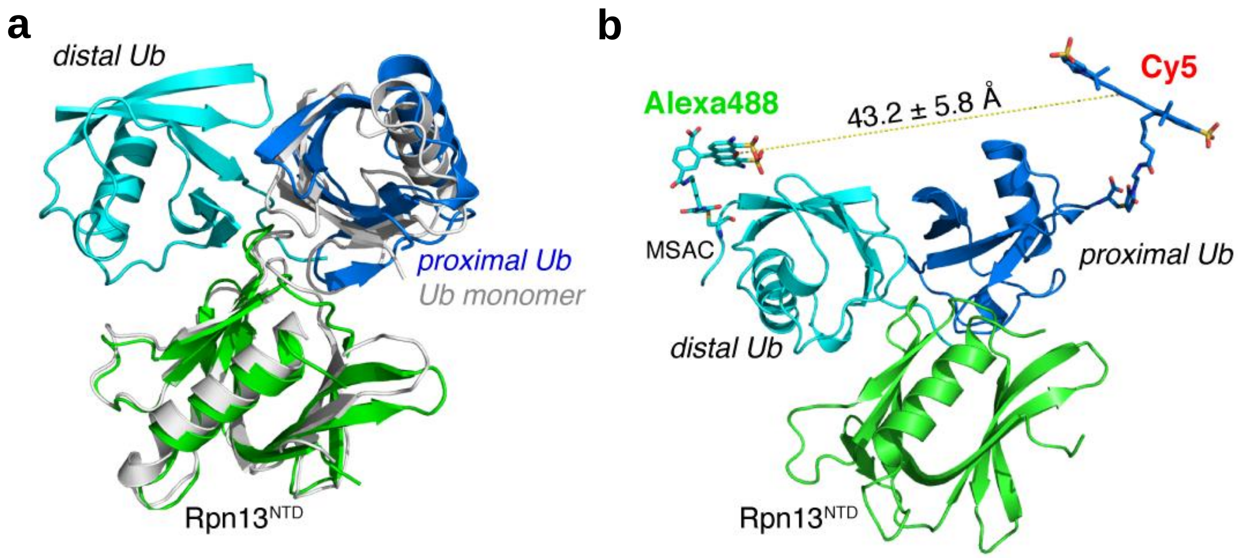

**Supplementary Fig. S10** Assessment of the complex structure between Rpn13$^{NTD}$ and K48-diUb. (**a**) The proximal Ub and Rpn13$^{NTD}$ in the complex is similar to the complex of Rpn13$^{NTD}$ and Ub monomer (gray cartoon, PDB code 2Z59). When superimposed by Rpn13$^{NTD}$ and proximal Ub, the backbone RMS difference is 2.17±0.31 Å. (**b**) FRET distance can be calculated between maleimide Alexa Fluor 488 modeled at the N-terminus of the distal Ub (0C, with residues MSAC appended) and maleimide Cy5 modeled at the C-terminus of the proximal Ub (76C). The averaged distance is 43.2±5.8 Å, corresponding to an average calculated FRET efficiency of 0.73.

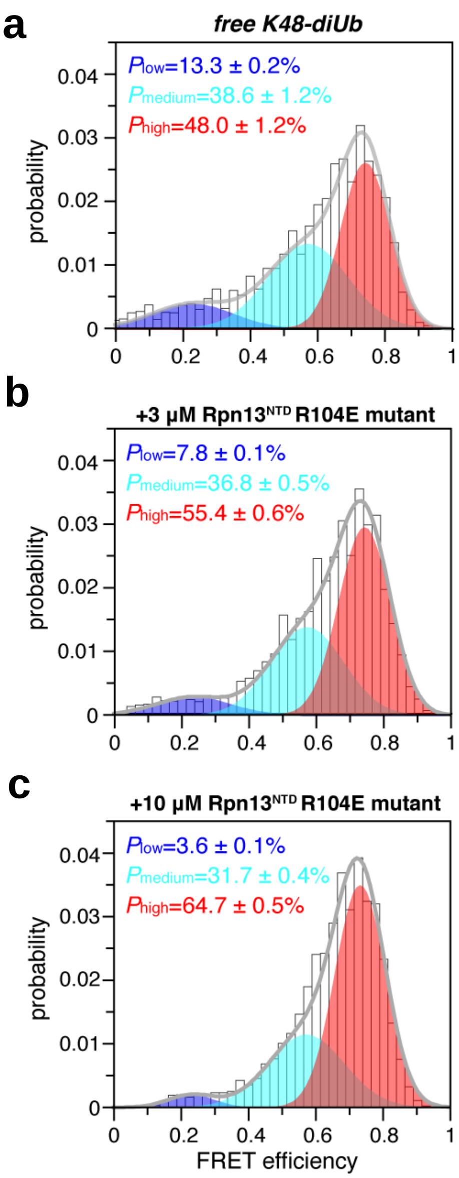

**Supplementary Fig. S11** The smFRET titration of K48-diUb with Rpn13$^{NTD}$ R104E mutant. Fluorophore-labeled K48-diUb (at 76C site of the proximal Ub and 0C of the distal Ub, Supplementary Fig. S1a) is titrated with Rpn13$^{NTD}$ mutant, carry a charge reversal mutation R104E. At 3 μM and 10 μM, Rpn13 mutant selectively enriches the population of the high FRET species by ~7% and ~17%, respectively.

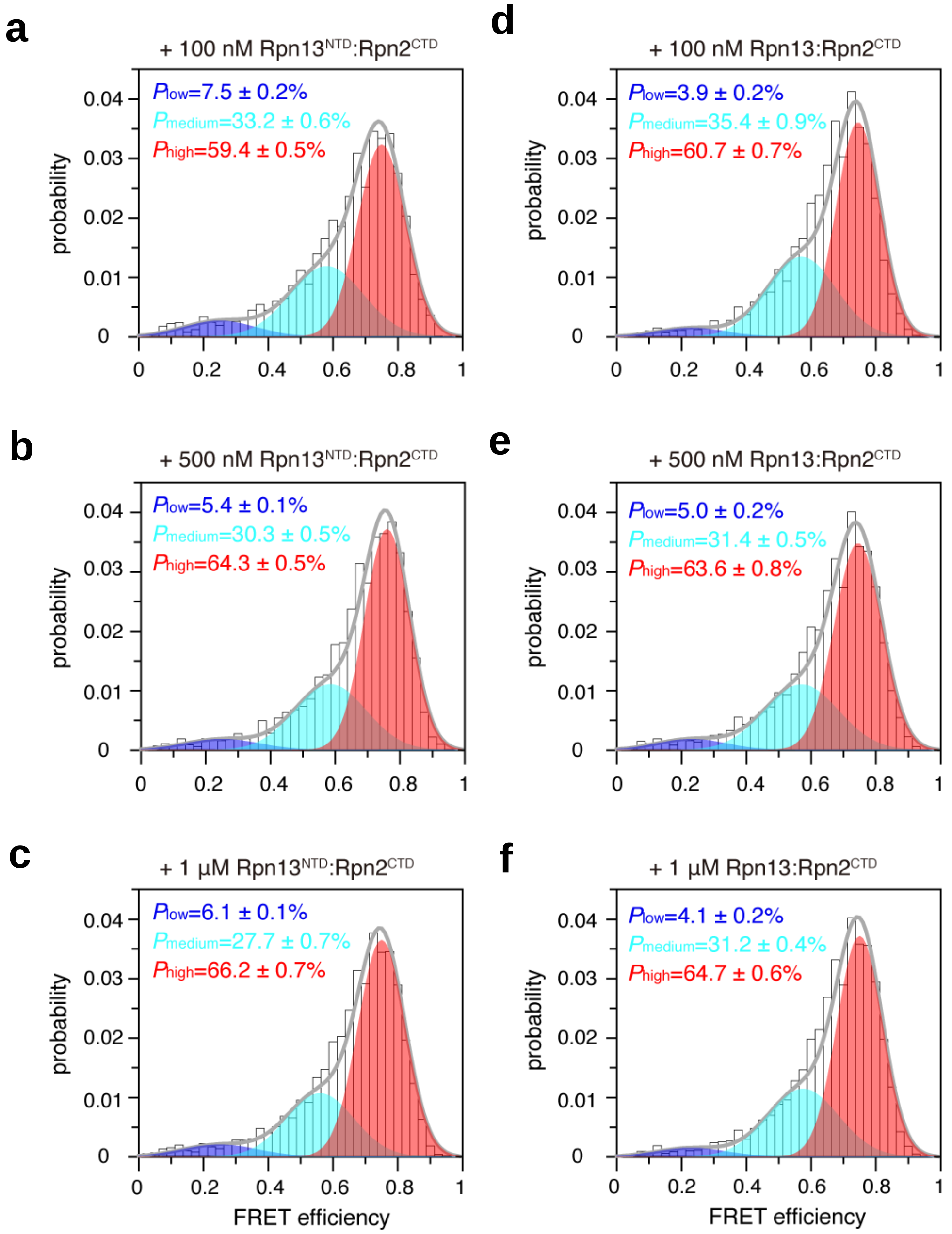

**Supplementary Fig. S12** The smFRET titration of fluorophore-labeled K48-diUb with 1:1 mixture Rpn13$^{NTD}$:Rpn2$^{CTD}$ or Rpn13:Rpn2$^{CTD}$. The fluorophores are labeled at 76C site of the proximal Ub and 0C of the distal Ub (Supplementary Fig. S1a). Both Rpn13$^{NTD}$:Rpn2$^{CTD}$ and Rpn13:Rpn2$^{CTD}$ enriches the high-FRET species with very similar trend. Averaged populations and standard deviations were obtained from three independent measurements of smFRET time-traces.

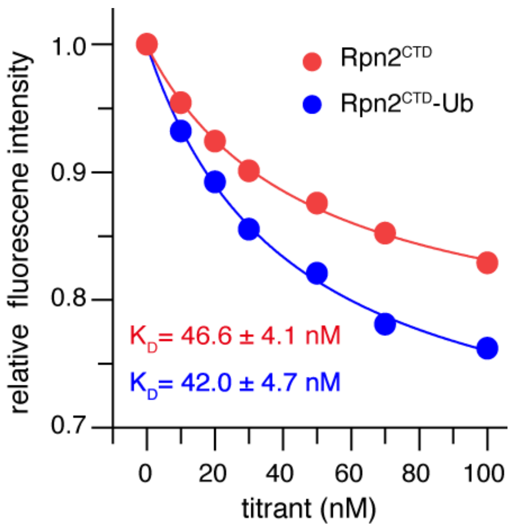

**Supplementary Fig. S13** Rpn2$^{CTD}$-Ub binds to Rpn13 with sub-μM affinity. Attachment of Ub at the C-terminus of Rpn2$^{CTD}$ does not affect the binding affinity between Rpn2$^{CTD}$ and Rpn13$^{NTD}$. The $K_D$ values were measured by evaluating the intrinsic Tryptophan fluorescence, as Rpn13 residue W108 is located at the interface with Rpn2.

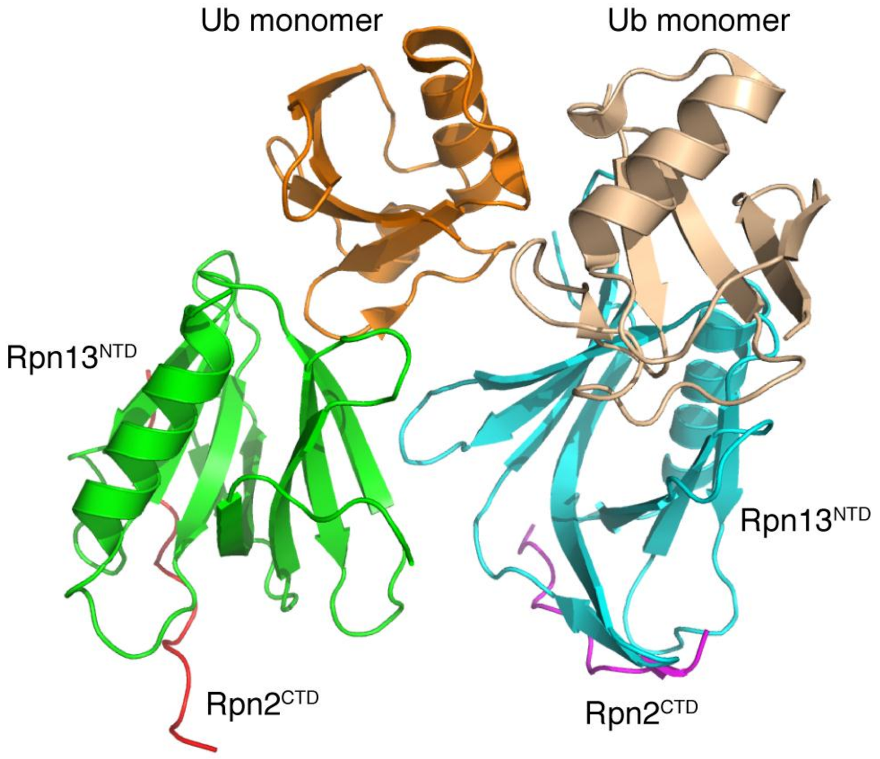

**Supplementary Fig. S14** The structure of Rpn13$^{NTD}$:Ub monomer has been crystalized as a dimer in one asymmetric unit (PDB accession code 5V1Y).

**Table S1** Structure statistics of Rpn13NTD:K48-diub complex

| | Rpn13$^{NTD}$:K48-diUb complex |
|---|---|
| **Structure bundle Size** | 20 |
| **Restraint statics** | |
| NOE RMS (Å) | 0.42±0.05 |
| PRE Q-factor | 0.90±0.003* |
| **Average Pairwise r.m.s deviation (Å)** | |
| all atoms | 1.40±0.31 |
| back-bone atoms | 0.98±0.39 |
| **Buried interface area (Å$^2$)** | |
| Ub:Ub | 1122.875±116.224 |
| distal Ub:Rpn13 | 941.010±79.048 |
| proximal Ub-Rpn13 | 1296.89±109.059 |
| **Z scores** | |
| 2nd generation packing quality | -1.584 |
| Ramachandran plot appearance | 0.484 |
| Backbone conformation | 1.173 |
| **Ramachandran scores** | |
| favorable | 92.6 % |
| additional allowed | 6.6 % |
| generously allowed | 0.8 % |
| disallowed | 0.0 % |

*The PRE restraints were applied as a square-well potential. Some residues experience large PREs and their peaks are completely broadened out in the paramagnetic spectrum (Fig. 3d, e). An arbitrary large PRE target value (> 120 s-1) was used; if excluding these residues, the PRE Q-factor for the final structures is 0.22±0.03.
